# Robustness of individualized inferences from longitudinal resting state dynamics

**DOI:** 10.1101/2020.09.15.297572

**Authors:** Maximilian Hommelsen, Shivakumar Viswanathan, Silvia Daun

**Affiliations:** 1Cognitive Neuroscience, Institute of Neuroscience and Medicine (INM-3), Forschungszentrum Jülich, Jülich, Germany; Institute of Zoology, University of Cologne, Cologne, Germany

**Keywords:** Neural plasticity, Individual differences, Individual identification, Electroencephalography (EEG), Power Spectrum, Frequency analysis, Machine learning, Multiclass classification

## Abstract

Tracking how individual human brains change over extended timescales is crucial in scenarios ranging from healthy aging to stroke recovery. Tracking these neuroplastic changes with resting state (RS) activity is a promising but poorly understood possibility. It remains unresolved whether a person’s RS activity over time can be reliably decoded to distinguish neurophysiological changes from confounding differences in cognitive state during rest. Here, we assessed whether this confounding can be minimized by tracking the configuration of an individual’s RS activity that is shaped by their distinctive neurophysiology rather than cognitive state. Using EEG, individual RS activity was acquired over five consecutive days along with activity in tasks that were devised to simulate the confounding effects of inter-day cognitive variation. As inter-individual differences are shaped by neurophysiological differences, the inter-individual differences in RS activity on one day were analyzed (using machine learning) to identify a distinctive configuration in each individual’s RS activity. Using this configuration as a classifier-rule, an individual could be re-identified with high accuracy from 2-second samples of the instantaneous oscillatory power acquired on a different day both from RS and confounded-RS. Importantly, the high accuracy of cross-day classification was achieved only with classifiers that combined information from multiple frequency bands at channels across the scalp (with a concentration at characteristic fronto-central and occipital zones). These findings support the suitability of longitudinal RS to support robust individualized inferences about neurophysiological change in health and disease.

## 1. INTRODUCTION

Tracking how individual human brains change over extended time-scales (e.g., days to years) is crucial to monitor and modify neural plasticity processes in scenarios ranging from healthy aging (Boersma et al. 2011; Cabeza et al. 2018; Cassani et al. 2018) to stroke recovery (Giaquinto et al. 1994; Rehme et al. 2011; Wu et al. 2016; Bonkhoff et al. 2020; Saes et al. 2020; van der Vliet et al. 2020). A promising strategy to track an individual’s changing neurophysiology is with repeated measurements of resting state (RS) activity, i.e., the ongoing neural oscillatory dynamics over a few minutes of wakeful rest (Vecchio et al. 2013; Carino-Escobar et al. 2019; Newbold et al. 2020; Pritschet et al. 2020; Saes et al. 2020). RS-activity has been shown to provide reliable indicators of neurobiological organization and integrity (Biswal et al. 1995; Damoiseaux and Greicius 2009; Van Den Heuvel et al. 2009; Hermundstad et al. 2013; Miŝic et al. 2016; Hoenig et al. 2018; Buckner and DiNicola 2019). The apparent informativeness of RS-activity as well as its convenient acquisition at relatively low cost (for example, with electroencephalography (EEG)) supports its relevance for long-term tracking. However, the relationship between RS <u>changes</u> over repeated measurements to neurophysiological <u>change</u> is poorly understood. Decoding this relationship is crucial to draw inferences about a person’s changing brain using longitudinal RS.

A basic inference required from longitudinal RS is about the origin of inter-day RS differences. Suppose a person’s RS-activity patterns *A_p_* and *A_q_* (on days *p* and *q*) are different. Is this difference attributable to (i) a possible neurophysiological change (abbreviated as *NP+*), or (ii) an incidental difference in inter-day activity (i.e., *NP-*)? Although an *NP+/NP-* decision involves many considerations, a key question is whether this decision is decodable from the relationship between *A_p_* and *A_q_*.

A major difficulty in decoding an *NP+/NP-* decision from RS-activity is the unconstrained format of the rest task. The rest task is defined by: (i) a *behavioral* state specified by instructions to stay still and keep eyes open (or closed) (Barry et al. 2007); and (ii) a *cognitive* state typically specified by instructions to relax and avoiding thinking of anything specific. Unlike the behavioral state, the criteria to objectively verify the cognitive state are ill defined (Benjamin et al. 2010; Duncan and Northoff 2013; Kawagoe et al. 2018). Due to this ambiguity, inter-day RS changes do not have a simple correspondence to neuroplastic change. For instance, a person’s incidental cognitive state during the rest-task could vary between days (e.g., session 1: free mind-wandering, session 2: struggling to stay awake, session 3: replaying emotional memories) (Diaz et al. 2013; Gonzalez-Castillo et al. 2021). The neural processing related to these differing cognitive states could in turn modify RS-activity *without* any changes to underlying neurophysiology. Therefore, large inter-day changes in RS-activity might not imply *NP+* and small changes might not imply *NP-.* Given this confounding potential built into the rest task, in the current study, we investigated whether RS-activity has other properties to support *NP+/NP-* classification.

Although inter-day RS *differences* are vulnerable to confounding by variable cognitive states, this might not be so for inter-day RS *commonalities*. We pursued this possibility by adopting a simple model of how inter-day RS commonalities might be structured. An individual’s neurophysiology on a particular day is assumed to impose constraints on how RS-activity is configured irrespective of cognitive state. This constraint-defined configuration would be shared by RS-activity across days only if these unique constraints are also shared. Such a configuration, if it indeed exists, provides a decision-rule for *NP+/NP-* classification as follows.

Suppose **C***_p_* denotes the constraint-defined configuration in the activity pattern *A_p_.* If activity *A_q_* on a different day is consistent with **C***_p_* then it supports an *NP-* classification, as inter-day consistency is assumed to require shared neurophysiological constraints. Conversely, if *A_q_* is not consistent with **C***_p_* then it suggests a change in these constraints and supports an *NP+* classification. As this constraint-defined configuration is assumed to be independent of cognitive state, the *NP+/NP-* decisions with such a decision-rule should presumably escape confounding by inter-day cognitive variability. Thus, according to this model, *NP+* and *NP-* are hypothesized to have distinctive, decodable signatures in RS-activity. Here, we sought to empirically test this predicted decodability of longitudinal RS.

Using EEG, longitudinal RS-activity was acquired on five consecutive days from a group of healthy, young participants. For these data, we sought to obtain decision-rules capable of *NP+/NP-* classification from the power spectrum of brief two-second samples of oscillatory activity at channels across the scalp. Testing the decision-rule’s robustness to confounding needed suitable samples known to be (i) free of neurophysiological change (*NP-*) and (ii) samples containing such changes (*NP+*).

Testing *NP-* decisions posed an experimental challenge. Our participants were assumed to be neurophysiologically stable across the five-day measurement period. However, the variation in their inter-day cognitive states during the rest-task was unverifiable. Therefore, the measurement protocol included two additional tasks to produce *pseudo-*rest states that were matched to rest in behavior but not in cognitive state (Figure 1A). These pseudo-rest states served to simulate confounding RS differences of varying sizes and complexity and allowed a rigorous test of the robustness of *NP-* decisions.

**Figure 1:**
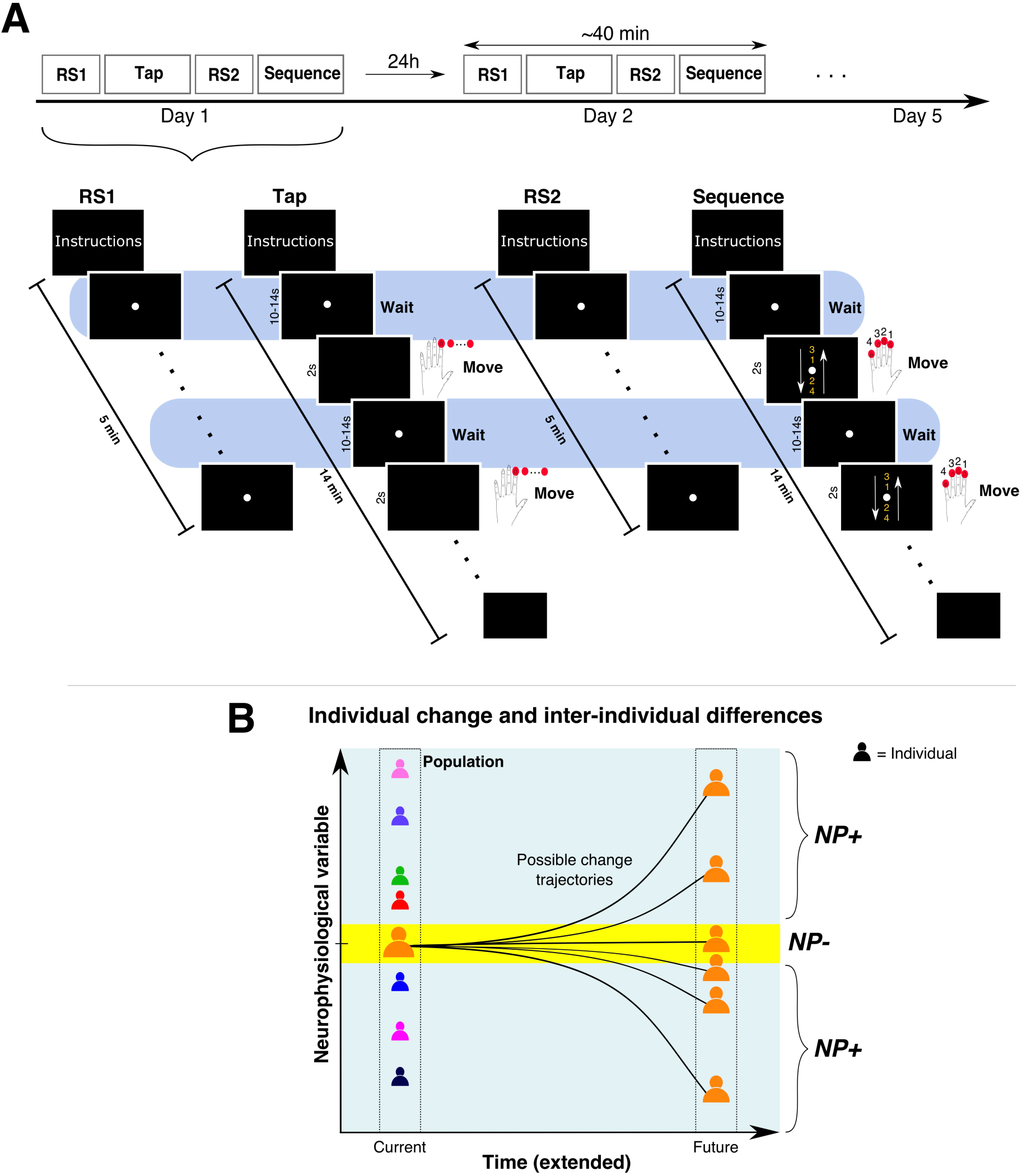
Experiment rationale. **(A)** Four tasks (*RS1, Tap, RS2, Sequence*) were performed in the same fixed order daily on five consecutive days. Task details for one day are schematically illustrated. A white fixation point was continuously displayed during the *RS1* and *RS2* task periods, and during “waiting” periods in the *Tap* and *Sequence* tasks (highlighted in blue). In the *Tap* task, a blank screen cued a 2s movement interval requiring left index-finger movements to repeatedly press a button (shown as red dots). In the *Sequence* task, the movement cue was an image depicting four numbers between two arrows (not drawn to scale) indicating the sequence of buttons to be pressed in a continuous cyclical manner, e.g., 3-1-2-4-4-2-1-3-3-1-2-4, etc. Number-to-finger mapping is shown on cartoon hand. **(B)** Schematic of longitudinal changes to a single hypothetical neurophysiological variable for one selected individual (orange). The current value of this variable (yellow area; *NP-*) can change in a variety of possible ways over long time-scales (gray area; *NP+*). The values of this variable in other individuals in a population cross-section (colored icons) provide proxies for these unknown individual-specific change trajectories.

Testing *NP+* decisions presented a methodological conundrum. Over long time-scales, an individual’s neurobiology could change in variety of possible ways (Cabeza et al. 2018; Grefkes and Fink 2020), with very different associated consequences for RS-activity (Figure 1B). These hypothetically possible RS-activity patterns were, by definition, experimentally inaccessible and limited the options for an individual-specific test of *NP+* classification. As a pragmatic alternative, we used a cross-sectional approach where RS-activity from *other* individuals served as simulated examples of RS-activity requiring an *NP+* classification. We assume that an individual *S*’s neurophysiology differed from other individuals in the tested population to variable extents. Therefore, relative to each individual *S,* the RS-activity of others provided a diverse range of examples of RS-activity with an origin in true neurophysiological differences.

By adopting the above strategy to obtain suitable *NP-* and *NP+* examples, the demands for *NP+*/*NP-* classification shared similarities to the demands for *person* identification, namely, obtaining a decision-rule to distinguish *S* from other individuals based on RS-activity (Figure 1B). Numerous prior studies demonstrate that RS-activity can serve as a “fingerprint” for person identification (Huang et al. 2012; Campisi and Rocca 2014; Finn et al. 2015; Valizadeh et al. 2019; Pani et al. 2020). Although our focus was not on the neural basis of individual differences and trait-identification (Smit et al. 2005, 2006; Demuru et al. 2017; Finn et al. 2017; Gratton et al. 2018), this person identification approach provided a convenient technical platform for our test of individual-specific longitudinal inference. Therefore, we mapped our test of robust cross-day *NP+/NP-* classification into the terminology of a person identification problem and adopted a machine-learning approach to address this problem.

Decision-rules (i.e., classifiers) were trained to distinguish a person *S* from all others in the tested population using samples from a single day. The samples from *S* putatively share a constraint-defined configuration that is not shared by samples from other individuals. Therefore, the outcome of training should be a decision-rule that represents information about individual S’s unique configuration. If this is indeed true, *S*’s decision-rule from one day should enable *S* to be re-identified from samples acquired on a different day as well as from samples of pseudo-RS activity despite cognitive state variability (*NP-*). Conversely, *S*’s decision-rule should classify samples from other individuals as not-*S*, consistent with a difference in neurophysiology co-mingled with cognitive state differences (*NP+*).

## 2. MATERIALS & METHODS

### 2.1. Participants

Twenty seven healthy volunteers (11 female, age (mean ± sd): 27.9 years ± 3.4, range: 22-34 years) participated in the study and received monetary compensation. Participants had normal or corrected-to-normal vision; no history of neurological or psychiatric disease; were not under medication at that time; and had no cranial metallic implants (including cochlear implants). Handedness was not an inclusion criterion. Based on the Edinburgh Handedness Inventory (Oldfield 1971), 22 participants were right handed (score > 50), 2 were left handed (score < −50) and 3 had intermediate scores. The study was approved by the Ethics Commission of the Faculty of Medicine, University of Cologne (Zeichen: 14-006). All participants provided their written informed consent before the start of the experiment.

Datasets from 24 (of the 27) participants were used for statistical analyses (see section 2.6).

### 2.2. Apparatus and EEG data acquisition

Stimuli were displayed using the software Presentation (v. 20.2 Build 07.25.18, Neurobehehavioral Systems, Inc.) on an LCD screen (Hanns-G HS233H3B, 23-inch, resolution: 1920 x 1080 pixels). Behavioral responses were recorded with the fMRI Button Pad (1-Hand) System (LXPAD-1×5-10M, NAtA Technologies, Canada).

Scalp-EEG was acquired with a 64-channel active Ag/AgCl electrode system (actiCap, Brain Products, Germany) having a standard 10-20 spherical array layout (ground electrode at AFz, reference electrode on the left mastoid). Three electrodes (FT9, FT10, TP10) were used to record electrooculographic (EOG) activity: one below the left eye to record vertical movements and the other two near the left and right lateral canthi to record horizontal movements. During acquisition, measured voltages (0.1µV resolution) were amplified by a BrainAmp DC amplifier (BrainProducts GmbH, Germany) at a sampling rate of 2.5 kHz and filtered (low cutoff: DC, high cutoff: 250 Hz).

To ensure reliable positioning of the EEG cap across sessions, a stereotactic neuronavigation system (Brainsight v. 2.3, Rogue Research Inc, Canada) was used on each session to co-register the spatial coordinates of five selected electrodes (AFz, Cz, POz, C5, C6) to their coordinates on the first session (see section 2.4 for details).

### 2.3. Experiment protocol and paradigm

Participant completed five sessions of approximately 40 minutes each (Figure 1A, upper panel) scheduled at the same time on consecutive days (Monday to Friday). Sessions took place at three possible times: morning (6 x 9AM), noon (9 x 12PM) and afternoon (12 x 3PM). Due to technical problems during the scheduled recording, for one participant, the fifth session was re-acquired after a gap of three days. For all participants, every session consisted of two resting state recordings (*RS1* and *RS2*) interleaved with two non-rest tasks (referred to as *Tap* and *Sequence*) in the same order (namely, *RS1*, *Tap, RS2, Sequence*).

The *Tap* and *Sequence* tasks (Figure 1A, lower panel) involved some special design considerations. Both tasks required participants to press buttons in response to visual cues. However, these tasks had relatively long and variable inter-stimulus-intervals (10-14s) where participants fixated on the screen as they “waited” for the visual cue that required the instructed response. The cognitive states during these waiting periods (referred to as *TapWait* and *SeqWait*) were the primary focus of these tasks. The behavioral demands of the *Tap* and *Sequence* tasks were designed to modulate the cognitive states during these pre-movement wait periods, for example, covert movement preparation during *TapWait* and covert rehearsal of a movement sequence during *SeqWait.* With this covert modulation, the *TapWait* and *SeqWait* could be considered <u>pseudo*-*rest</u> states as they were matched to *RS1* and *RS2* in behavioral state but not in cognitive state. Furthermore, the *Tap* task was intended to produce cognitive states that were similar within and between days while the *Sequence* task was designed to elicit cognitive states that could systematically change across days. This was implemented by inducing participants to learn a difficult motor sequence where performance could improve with increasing practice across days. We now describe the different task periods in detail.

Each task period began with an instruction screen describing the task to be performed and ended with another instruction screen that instructed participants to take a short break and press a button to initiate the next part when they were ready.

#### 1: Resting State (RS1)

During this period lasting ∼5minutes, a small white dot was continuously displayed at the center of the screen. Participants were instructed to keep their eyes open, fixate on the displayed white dot, relax and avoid movements (also see section: Procedure).

#### 2: Tap task

In this task-period, a small white dot was centrally displayed on the screen (as in *RS1*).

However, after variable intervals of 10-14 seconds, this dot disappeared for a 2 second period before reappearing. The offset of the dot was the cue for participants to repeatedly and rapidly press a button with their left index finger until the dot reappeared on the screen. The total task (duration ∼14 minutes) consisted of 60 movement periods (dot absent) interleaved with 60 waiting periods (dot present). These waiting periods are referred to as *TapWait* and the response execution periods are referred to as *TapMov*.

#### 3: Resting State (RS2)

A second resting state recording was acquired with all task parameters being identical to *RS1*. This recording is referred to as *RS2*.

#### 4: Sequence task

As with the *Tap* task, the sequence task consisted of 60 waiting periods of 10-14s each (i.e., *SeqWait*) where a small white dot was centrally displayed on the screen interleaved with 60 movement periods of 2s duration (i.e., *SeqMov*). Unlike the *Tap* task, each movement period was cued by a centrally displayed visual stimulus consisting of four vertically displayed digits (3-1-2-4) between two vertical arrows. Each number was mapped to a different button on the response pad. The vertical ordering of the numbers indicated the sequence in which the indicated buttons had to be pressed using fingers of the left hand. The arrows indicated that this sequence had to be executed rapidly and repeatedly in a cyclical manner starting from top to bottom and back. For example, following stimulus onset, the required sequence of button-presses was 3-1-2-4-4-2-1-3-3-1-2-4-… and so on. This continuing sequence had to be executed until the offset of the stimulus. No performance feedback was provided during the task. This particular sequence of digits was selected as it was challenging to execute rapidly. To promote learning of this sequence across trials and days, the same sequence of numbers and number-to-finger-mapping was used on all sessions. The same sequence and number-to-finger mapping was also used for all participants.

Handedness was not an inclusion criterion in our experiment. However, for uniformity in task-related neural activity, all participants used fingers of their left hand to execute the button-press responses in the *Tap* and *Sequence* tasks.

### 2.4. Procedure

Prior to the start of the recordings on each of the five days, participants completed the Positive and Negative Affect Schedule (PANAS) (Watson et al. 1988) and completed brief questionnaires to report the caffeine consumption on that day and the amount and quality of sleep on the previous night.

On the first day, participants received detailed instructions about the experiment. For the resting state periods, participants were instructed to keep their eyes open, fixate on the displayed white dot and to avoid movements. Additionally, they were also asked to relax, stay awake and not think of anything in particular. For the *Tap* task, participants were instructed to press the buttons as rapidly as possible without causing discomfort. For the *Sequence* task, participants were familiarized with the task and the mapping of the number to finger. They practiced performing the task using a different digit sequence from the one used in the main experiment. Furthermore, they were explicitly instructed on each session to try to improve their performance particularly the number of buttons pressed during each response period. Finally, on all sessions, we repeatedly emphasized the importance of minimizing eye-blinks, maintaining fixation at all times during the recording, and the avoidance of all unnecessary movements of the fingers, head and body.

As the study’s objective was to relate the spatio-temporal organization of neural activity across days, minimizing inter-day variation in the EEG cap’s position was an important priority. We therefore implemented an additional spatial registration procedure on each day after the EEG cap was secured to the participant’s head. Using a stereotactic neuronavigation system, the participant’s head was registered to the Montreal Neurological Institute (MNI) space using standard cranial landmarks. The positions of five selected electrodes along the midline and lateral axis (AFz, Cz, POz, C5, C6) were then localized using the neuronavigation software. The electrode locations obtained on the first day were used as the reference for the remaining sessions. On each subsequent session, the positioning of the cap was interactively adjusted so that each electrode’s coordinates closely matched its reference location. Due to scheduling constraints, this spatial registration procedure was not performed for 7 participants.

The application of electrode gel followed after cap positioning. Skin-electrode impedance was brought below 10kΩ before starting the recording. Recordings were acquired in a light-dimmed and acoustically shielded EEG chamber. Participants were seated in a comfortable chair with their heads stabilized with a chinrest in front of the computer screen at a viewing distance of ∼65cm. The response pad was placed in a recess under the table so that participants could not see their hands during the task-periods especially while pressing the buttons. During the recording, participants were monitored via a video camera to ensure that they maintained fixation, minimized eye-blinks, and stayed awake.

### 2.5. EEG preprocessing

The EEG data were preprocessed using the EEGLAB software (Delorme and Makeig 2004) and custom scripts in a MATLAB environment (R2016b, MathWorks, Inc., Natick, MA).

The continuous recordings were down-sampled to 128Hz, and then band-pass filtered to the range 1Hz-40Hz with a Hamming windowed sinc FIR filter (high pass followed by low pass). The continuous recordings then underwent an artifact correction process to remove oculomotor activity related to eye-blinks and saccades.

Eye blink removal was performed separately for each day’s dataset (including all task periods) using the procedure described by Winkler et al. (2015). Following this procedure, a copy of a dataset was first created which was then filtered with a high-pass 2 Hz filter. This duplicate dataset was visually inspected to remove data segments and EEG channels with artifacts related to repeated paroxysmal amplitudes changes (> 50µV), electromyographic contamination, electrical noise and signal loss. Next, the artifact-free data from all task-periods were segmented into 2s epochs. These epochs were then submitted to an Independent Components Analysis (ICA) decomposition using the infomax-ICA algorithm (implemented as *runica* in EEGLAB). To minimize inter-condition biases, ICA was performed on a balanced mixture of epochs from *RS1, TapWait, RS2* and *SeqWai*t. Epochs from the *TapMov* and *SeqMov* periods were excluded from this step to avoid movement-specific biases. The ICA weights obtained with the duplicate dataset were then transferred and applied to the original, non-filtered dataset. ICA components related to eye-blinks and saccades were then identified and removed using an automatic detection algorithm ADJUST (Mognon et al. 2011).

Following eye-blink correction, the original dataset was then again visually inspected to remove time periods and channels with artifacts. The signals in rejected channels were replaced with signals interpolated from other channels using spherical spline interpolation. All channels were then re-referenced to the Common Average Reference. Finally, the visually inspected continuous data were segmented into 2s epochs according to the six different experimental states: *RS1*, *RS2*, *TapWait*, *TapMov*, *SeqWait* and *SeqMov.* The epoch duration of 2s was heuristically selected to meet the tradeoff of (i) being short enough to obtain a sufficient number of samples for the machine-learning analysis (see section 2.6) while (ii) being long enough to obtain a suitable estimate of the power spectrum. Furthermore, this allowed epochs from the non-movement periods to match the 2s duration of the task-defined movement period.

For the two movement-related states (*TapMov* and *SeqMov),* epochs were defined from +0.25s to +2.25s following the visual cue to exclude initial transients and response-time delays following cue onset and to include residual movements in the period immediately following the cue offset. To avoid any carry-over effects from movement into the *TapWait* and *SeqWait* epochs, a time interval of 500ms immediately prior to cue onset and 1000ms immediately following cue offset were excluded before segmenting the *TapWait* and *SeqWait* epochs. Furthermore, all *TapWait* and *SeqWait* epochs that contained button presses were excluded.

To establish face-validity of the task states based on their time-courses, we created a separate set of epochs from −1 to +3s relative to the onset of the visual cue. The signals were band-pass filtered in the *β* frequency band (14-30Hz) and the signal amplitude was extracted using the Hilbert transform. After removing edge artifacts, the signal was normalized by calculating the percentage change in the signal relative to the mean amplitude in the pre-stimulus period [-898ms, 0ms]. After normalization, the signals were averaged across epochs, days and individuals.

### 2.6. Data quality assessment

Preprocessing resulted in 135 datasets (27 participants x 5 days). To be included in our analysis, each subject had to have completed the first three of the four tasks on all sessions and have at least 4 (out of 5) session-datasets that met the following data-quality criteria for analysis. We required a preprocessed dataset to have (i) less than seven rejected channels, (ii) ≥ 90 artifact-free epochs from both resting state periods (i.e., *RS1* and *RS2*), and (iii) ≥ 90 artifact-free epochs from the available resting-matched conditions (i.e., *TapWait, SeqWait*). Note that the number of epochs for *TapMov* and *SeqMov* were necessarily ≤ 60 as each task only had 60 response periods of 2s duration.

Datasets from 24 out of 27 participants met these data-quality criteria: 18 (of 24) had completed all 4 task-periods on each session and the remaining 6 (of the 24) participants had completed only the first 3 (of the 4 parts). To maintain uniformity in the statistical analyses, final analyses were performed only on the best 4 of the 5 session-datasets. For participants where all 5 datasets were of high quality, we excluded the first day’s dataset as it might involve effects of initial familiarization. To maximize the use of the available data after these exclusions, analyses involving only *RS1* and *RS2* included data from 24 participants, while analyses involving any of the non-rest tasks used data from 18 participants. For these 18 participants, the mean number of epochs per day in *TapMov* was 52.7 (min: 45.3, max: 57.7; SD = 2.9; minimum/day = 36) and in *SeqMov* was 53.4 (min: 49.2, max: 57.7; SD = 2.5; minimum/day = 42).

### 2.7. Feature specification: Oscillatory power spectrum

All classification analyses were based on a description of the oscillatory power spectrum on each 2-second epoch. Each epoch’s power spectrum was described using 305 features that specified the power in five canonical frequency bands (δ: 1-3.5 Hz; θ: 4-7.5 Hz; α: 8-13.5 Hz; β_1_ (low β): 14-22.5; β_2_ (high β): 23-30 Hz) at each of the 61 channels.

These features were extracted with the procedure schematically displayed in Figure 2A. For each 2s epoch of EEG activity, the oscillatory power spectrum at each channel over the range of 1 to 30 Hz (0.5Hz resolution) was computed using the Fast Fourier Transform (FFT). The power at all frequencies within each band’s frequency range was averaged to obtain the mean power per frequency band. The mean power per band was then logarithmically transformed (base 10) so that the resulting distribution across epochs had an approximate normal distribution. These five features (one per band) provided a minimal description of each channel’s power spectrum. Finally, these five features from each channel were concatenated to obtain a single vector with 305 feature values (5 frequency bands x 61 channels). This extended feature set describing an epoch’s power spectrum across the scalp was used for the classification analyses.

**Figure 2:**
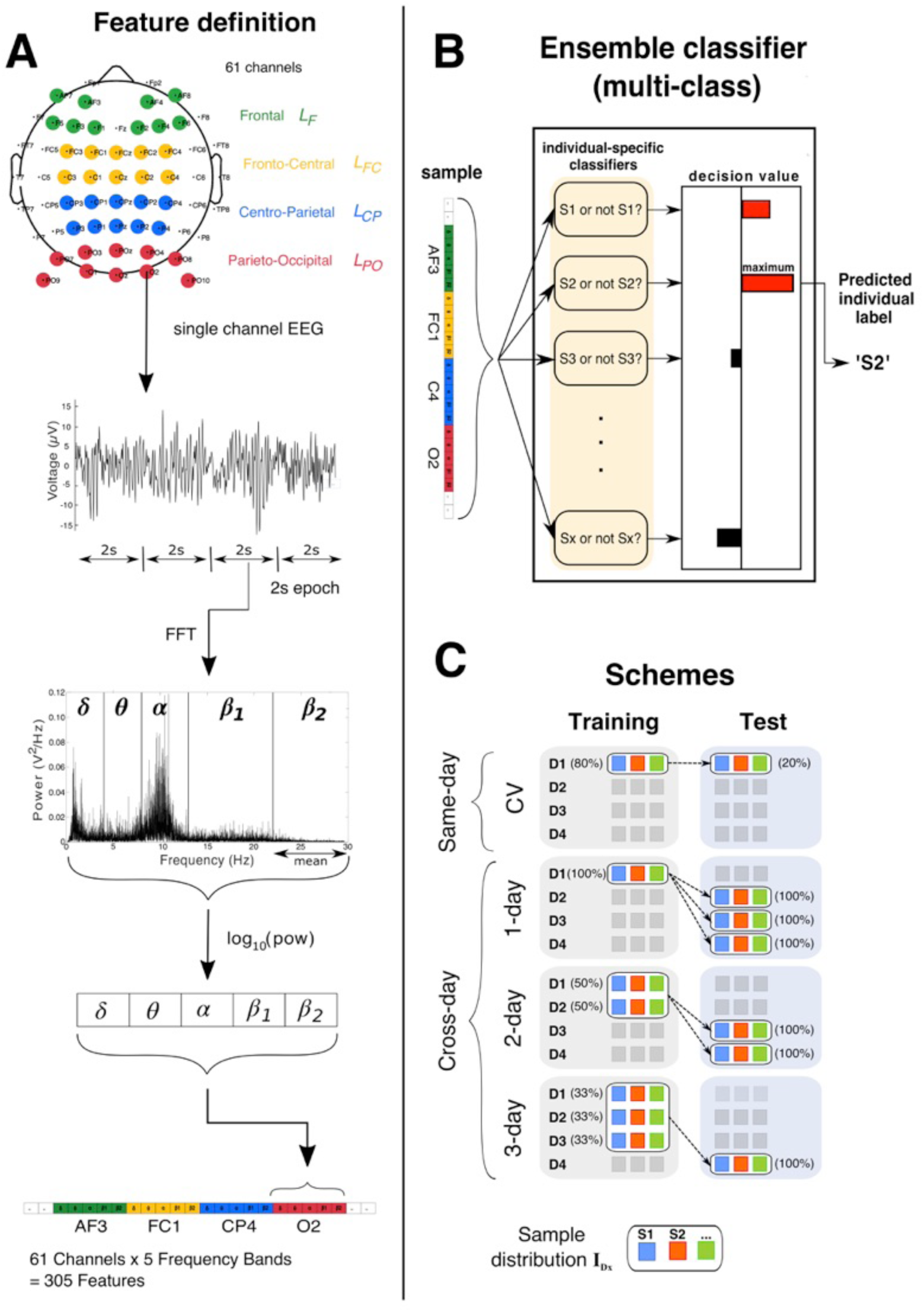
Classification procedure. **(A)** Feature definition pipeline. Channels in each mono-location subset are identified by color (green: *L*_F_, yellow: *L*_FC_, blue: *L*_CP_, red: *L*_PO_). The continuous signal from each channel was segmented into 2s epochs followed by an estimation of the frequency spectrum with the Fast Fourier Transform (FFT). The mean power within each of the five bands was log transformed (base 10) and concatenated with corresponding values from all other channels to obtain a feature vector. **(B)** Schematic of a multiclass decision with an ensemble of individual-specific binary classifiers. Each classifier evaluates the sample (*Sx* or not-*Sx*) to output a decision-value (red bars > 0, black bars < 0) and the classifier with the maximum decision value was the predicted label (here, S2). **(C)** Classification schemes ^A^**I**_Dx_ → ^A^**I**_Dy_ (rows) were defined by the configuration of training (left column) and test sets (right column) (where **Di** denotes samples from day *i*). The sample distribution (**I**_Dx_) had samples from all individuals (multi-colored boxes). Percentages indicate the proportion of each day’s samples used for training/testing. Same-day identification was estimated with 5-fold cross-validation (CV). The training set for cross-day aggregation had an equal proportion of samples from each day and the total number of training samples was the same across aggregation levels.

For detailed analyses, we defined subsets of the full feature set referred to here as the (i) mono-band and (ii) mono-location feature sets. Each *mono-band* feature set (*B_f_*) consisted of features belonging to only one frequency band *f*. The five mono-band feature sets (each with 61 features) were *B*_δ_, *B*_θ,_ *B*_α_, *B*_β1_ and *B*_β2_. Each *mono-location* feature set (*L_z_*) (Figure 2A, top panel) consisted of features from 10 bilaterally symmetric channels in the spatial zone *z* on the scalp along the anterior-posterior axis. The four mono-location sets were defined at the frontal (*L_F_*); fronto-central (*L_FC_*), centro-parietal (*L_CP_*) and parieto-occipital (*L_PO_*) zones respectively.

### 2.8. Multi-class classification

#### 2.8.1. Definition

All classification models were numerically estimated using a soft-margin linear Support Vector Machine (SVM, with *L2* regularization) algorithm as implemented by the *LinearSVC* package in the *scikit-learn* library (Pedregosa et al. 2011) implemented in Python 3.6. SVM learning was initialized with parameters (tolerance = 10^-5^, max iterations =10^4^, hinge loss, balanced class weighting). The hyper-parameter *C* had a value of 1, which has been shown to be a reasonable default for M/EEG classification (Varoquaux et al. 2017). For our data, tuning *C’*s value seemed to produce only marginal changes to the classification accuracies (results not shown).

As defined above, each epoch was a 2-second sample of the ongoing oscillatory activity from one person (of 24) on one specific day (of 4) engaged in a particular task state (of 6 possible states: true rest {*RS1*, *RS2*}, pseudo-rest {*TapWait, SeqWait*}, non-rest {*TapMov, SeqMov*}). The classification analyses involved predicting an epoch’s origin either by (i) a person’s identity or (ii) task-state. Multi-class classifiers (using an ensemble of binary classifiers) were used for person identification as described below. Standalone binary classifiers were used to distinguish alternative task-states within the same person.

The input to a multi-class classifier (see Figure 2B) was a single sample (i.e., epoch) from an unspecified person *S*_x_ in the studied group and the required output was the predicted identity of that person (e.g., *S_2_*). The multi-class classifiers used here employed a *one-vs-all* scheme (as implemented by *scikit-learn*). Specifically, an *N*-class classifier (*N* ≥ *2*) consisted of an ensemble of *N* binary-classifiers. Each of these binary classifiers was independently trained to distinguish whether a sample was from one specific person (e.g., *S*_2_) or from any of the other *N-1* persons (i.e., not *S*_2_). Therefore, each individual was associated with a unique classifier in the ensemble. To obtain a classification with such an ensemble, each sample was separately evaluated by each of the *N* binary-classifiers to obtain a decision value from each classifier (i.e., the signed distance to the separation hyperplane (Rifkin and Klautau 2004)). These decision values were compared and the final classification was assigned to the binary classifier with the maximum decision value.

#### 2.8.2. Accuracy scoring

Even though an ensemble was used for multi-class classification, our interest was in the accuracy of each individual-specific binary classifier in the ensemble. To obtain a measure of classification accuracy of each individual classifier from the ensemble classification accuracy, we defined the accuracy *a*_i_ of the classifier for person *S_i_* as

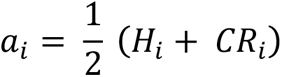

where *H_i_* denotes the hit rate (i.e., positive identification rate) of the classifier and *CR*_i_ denotes the correct rejection rate. The hit rate *H_i_* was the proportion of instances where samples from *S_i_* were correctly predicted as being from *S_i_* by the ensemble (i.e., a true positive where the classifier *S_i_* had a larger decision value than the competing classifiers). Correct rejection was defined based on the pair-wise relationship of *S_i_* to each of the other classifiers *S_j_.* If the ensemble (incorrectly) predicts *S_i_* for a sample from a different person *S_j_* then it implies that the classifier *S_i_* (incorrectly) had a larger decision value than the competing classifiers, i.e., a false positive. The false positive rate *FP_i,j_* denotes the proportion of instances where samples from *S_j_* were incorrectly predicted as being from *S_i_* by the ensemble. The correct rejection *CR_i,j_* was defined as *CR_i,j_ = 1 - FP_i,j_*. Based on this rationale, the overall correct rejection *CR_i_* for *S*_i_ was defined as the mean of the pair-wise correct rejection rates

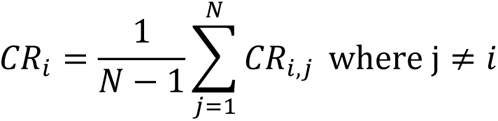

With this formulation, random chance for each classifier was 50% even though random chance for the entire ensemble was (100/*N)%*.

To identify individuals who were frequently misclassified (i.e., confused) with each other, we report confusion matrices for cross-day classification. In this confusion matrix, the rows represent the true label of a sample and the columns indicate the predicted label for that sample by the ensemble. The value for the row corresponding to individual *S_i_* and column corresponding to individual *S_j_* indicated the proportion of samples from *S_i_* that were classified as *S_j_*. The rows/columns of the matrices were re-organized to cluster together individuals who were confused with each other. This was implemented with the so-called Louvain method to maximize modularity (Blondel et al. 2008), implemented in the Community Detection Toolbox (Kehagias 2021).

The accuracy score can have different contributions from the hit-rate (e.g., high false negatives) and the correct rejection rate (e.g., high false positives). To disentangle these contributions, we estimated the *recall* and *precision* scores from the confusion matrix (Davis and Goadrich 2006). The *recall* score for individual *S_i_* is the ratio (True Positives)/(True Positives + False negatives). The recall score for *S_i_* would be low if samples from *S_i_* are misclassified as belonging to another individual (i.e., false negatives). The precision score for individual *S_i_* is the ratio (True Positives)/(True positives + False positives). The precision score for *S_i_* would be low if samples from other individuals are misclassified as belonging to *S_i_* (i.e., false positives).

#### 2.8.3. Training and testing schemes

Classification was defined by the samples used for training and testing. Irrespective of classifier type (multi-class or standalone binary classifier), the training data were always balanced, (i.e., having an equal number of samples per class) to avoid biases arising from imbalanced classes (Abraham and Elrahman 2013).

Person (multi-class) identification was organized into two schemes based on whether the training and test samples belonged to the (i) same day (namely, same-day vs cross-day identification) and the (ii) same task (namely, same-task vs cross-task identification). A schematic of the same-day/cross-day schemes are shown in (Figure 2C). For convenience, we use the following notational convention to describe these classification schemes. As multi-class classification involves an ensemble decision, it involves the conjoint influence of the sample distributions from multiple persons. This combined distribution on a particular state (e.g., *RS1*) on day *d* is denoted as ^RS1^**I**_d_. A classification scheme where a decision-rule is trained on samples from ^A^**I***_p_* (i.e., from task state *A* on day *p*) and tested on samples from ^B^**I***_q_* (i.e., from state *B* on day *q*) is denoted as ^A^**I***_p_* → ^B^**I***_q_*. Similarly, a classification scheme where a decision-rule was trained on samples aggregated from different days (e.g., ^A^**I***_p_* and ^A^**I***_q_*) and tested on ^B^**I***_r_*. is denoted as ^A^**I***_p_* ∘ ^A^**I***_q_* → ^B^**I***_r_*. (see below).

##### Same-day/same-task identification

The accuracy of same-day person identification in task state *A* (^A^**I***_p_* → ^A^**I***_p_*) was estimated using a 5-fold cross-validation (CV) procedure (Blum et al. 1999). Specifically, the set of samples from state *A* on one day (for example, day D1 in Figure 2C, upper row), were partitioned into 5 equal folds. Training was performed on four folds (80% of the sample set) and tested on the left-out fifth fold (the remaining 20%). This training-testing procedure was repeated so that each fold was used as a test-set once. The mean classification accuracy across folds was defined as the same-day identification accuracy for that day. In this manner, the CV accuracy was estimated separately for each of the four days and the mean CV accuracy across days was denoted as the same-day accuracy for task state *A*.

##### Cross-day/same-task identification

For cross-day identification in task state *A* (^A^**I***_p_* → ^A^**I***_q_*), samples in the test set were from a different day than the samples in the training set. We modulated the training set’s day-specificity by aggregating samples from different days in a stratified manner. In an *n*-day training set, the *k* training samples per person consisted of *k*/*n* samples from each of *n* different days. Here, *n* could take three possible values, namely, 1, 2 or 3 (see Figure 2C, first column). The number of samples per person, *k*, was held constant to enable comparison of classification accuracy across all values of *n*. Irrespective of the extent of aggregation in the training set, samples in the test-set were never aggregated from different days. Mean identification accuracy for a particular *n*-day aggregation scheme (e.g., ^A^**I***_d1_* ∘ ^A^**I***_d2_* … ^A^**I***_dn_* → ^A^**I***_r_*.) was obtained by (i) independently estimating the accuracy for each possible training/test-set combination that satisfied the day constraints (e.g., day *p* ≠ day *q* ≠ day *r*) and then (ii) averaging these accuracy values.

##### Cross-task identification

This was treated as a special instance of cross-day identification. For example, the accuracy for the configuration ^A^**I***_p_* → ^B^**I***_q_* was estimated by replacing the test set with samples from state *B* while retaining all other day-related constraints as in cross-day/same-task identification. Unless specified otherwise, cross-task identification was always tested across days, that is, the training and test sets were always from different days. This was done to exclude potential inter-state similarities that might be present due to the joint preprocessing of data from all states within the same day (see section 2.5).

#### 2.8.4. Classification schemes: Interpretation of accuracy relationships

The same-day accuracy for a particular state was treated as a pre-condition to estimate the cross-day identification accuracy for that state. If same-day accuracy were greater than random chance, it would confirm that the distribution from which the training set was drawn contained sufficient information to allow identification in the absence of potential inter-day changes. Cross-day accuracy is reported and interpreted here only if this pre-condition was satisfied.

Based on this pre-condition, a reduction in cross-day (1-day) accuracy (e.g., ^A^**I***_p_* → ^A^**I***_q_*) relative to same-day accuracy (e.g., ^A^**I***_p_* → ^A^**I***_p_*) can be attributed to a systematic difference in the distributions ^A^**I***_p_* and ^A^**I***_q_* between days (red arrow, Figure 3A). Aggregation was used to evaluate the source of this cross-day accuracy reduction by varying the statistical properties of the training set (i.e., by aggregating samples across days) while holding the properties of the test set constant. Specifically, we assumed aggregation would lead to decision-rules that discount day-specific properties in favor of day-general properties. Therefore, depending on the relative roles of day-specific/general properties in the classification decision, the cross-day accuracy might stay constant, increase or decrease with increasing aggregation (Figure 3A).

**Figure 3:**
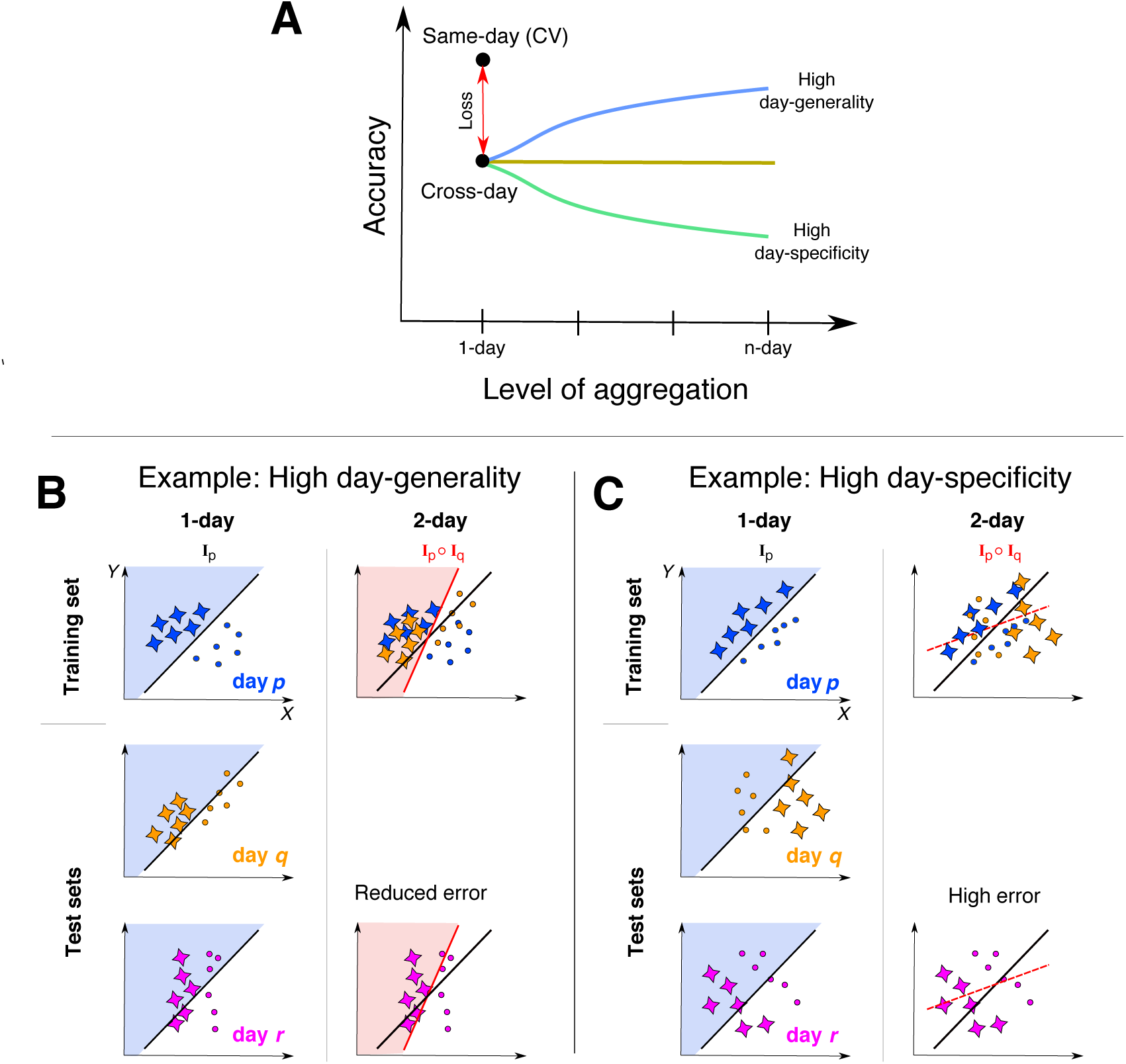
Effect of aggregation. **(A)** Schematic of relationship between same-day and cross-day identification accuracy. Cross-day (1-day) accuracy can be lower than same-day accuracy (red-arrow) due to day-specificity of the decision-rule. Training decision-rules on aggregated samples (y-axis) can change cross-day accuracy, which could increase (blue, see panel **B**), or stay constant (dark green), or even decrease (light green, see panel **C**). (**B**) Idealized example of how cross-day accuracy (column 1) can increase with aggregation (column 2) due to day-general information. Samples from two classes (stars, circles) are shown along two features (day-general: *X*, day-specific: *Y*) with each day’s samples shown in different colors (*p*: blue, *q*: orange, *r:* purple). The 1-day decision-rule (**I**_p_) (top left panel) is depicted with thick black line and shaded areas. This decision-rule can successfully classify samples from days *q* and *r* but with some errors. However, a decision-rule trained on data from days *p* and *q* (**I**_p_ ∘ **I**_q_) (thick red line, red shaded area) reduces cross-day classification errors (lower right). **(C)** Idealized example of high day-specificity. Even though the classes are separable within each day, the 1-day decision-rule (**I**_p_) has a poor cross-day accuracy (column 1). 2-day training (column 2) produces a decision-rule with worse classification both on the training set itself (dotted red line) as well as across days (lower right).

Figures 3B and 3C show idealized examples of how aggregation could both increase as well as decrease cross-day accuracy. In the example shown in Figure 3B, the two classes systematically differ on feature *X* (x-axis) but with an inconsistent role for feature *Y* (y-axis). Due to incidental day-specific variation, feature *Y* has a role in distinguishing the classes on day *p* but not on other days. Consequently, a decision-rule trained on day *p* does not effectively separate the classes on other days (column 1). However, training on aggregation samples from day *p* and *q* (column 2) reduces *Y*’s role in the aggregated decision-rule leading to an improved separation of the classes across days. Figure 3C illustrates an extreme example of day-specificity where the two features have a conjoint relationship allowing classification within each day but with low generality across days. Therefore, training on samples aggregated from day *p* and *q* leads to an overall reduction in accuracy on the training set itself as well across days.

#### 2.8.5. Weights and normalized weights

The characteristic weights for a particular classification scheme (e.g., ^A^**I***_p_* → ^A^**I***_q_*) were obtained by averaging the weights across all training sets. In a multiclass classifier, the decision-rules are organized in a winner-take-all competition to label each sample (Figure 2B). Therefore, for each sample to be uniquely assigned to only one person, the person-specific classifiers in the ensemble necessarily require different decision-rules. This difference in decision-rules might only be in the sign (positive/negative) assigned to the weights. Therefore, for all weight-related analyses, the absolute values of the weights were used in order to allow inter-individual comparisons.

To identify the high-consistency weights, the absolute weights were z-scored across all features for each subject to retain information about inter-feature weight differences in the statistical tests. However, this “raw” weight measure does not account for power differences. For features *i* and *j,* the weight |*w_i_|* might be greater than |*w_j_|* while the power |*P_i_*| might be less than |*P_j._*|. Consequently, neither the relationship between the weights nor the power are reliable indicators of the relative influence of *i* and *j* on the eventual classification decision. Therefore, we defined a feature *i*’s unit weight 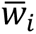 as the idealized weight value such that 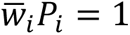. The normalized weight was thus defined as the ratio 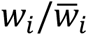, which was effectively equal to *w_i_P_i_*. Due to the characteristic differences in power between bands, for statistical comparisons, the absolute normalized weights (i.e., |w*P|) were z-scored <u>within</u> each band for each subject.

### 2.9. Statistical Analysis

The relative differences in the accuracy of different classification schemes were assessed by performing paired t-tests, repeated measures one-way or two-way analysis of variance (ANOVA) implemented by the pingouin python package (version 0.3.2) (Vallat 2018).

The random chance accuracy for the multi-class and standalone binary classifier was 50% and accuracy deviations from random chance were evaluated with one-sample t-tests. The Bonferroni threshold was used to correct for multiple comparisons. Due to the sequential relationship between the different multiclass classification schemes, following Figure 3A, the tests for same-day accuracy (CV) and cross-day accuracy were planned tests that were considered significant at a threshold of *p* < 0.05. The tests for 2-day and 3-day aggregation were evaluated at a threshold of *p* < (0.05/2). For tests for the mono-band and mono-location sets, the thresholds were further corrected for the number of feature sets. Correlations between individual accuracy values were evaluated using Spearman’s rank correlation due to the focus on relative ordering rather than a strict cardinal relationship.

Two kinds of error-bars are used in the plots. For plots depicting variable changes due to a single-factor, error bars indicate the standard deviation (SD). Plots depicting multi-factor changes use error bars displaying the within-subject standard error (s.e.m.) (O’Brien and Cousineau 2014). The type of error-bar used is explicitly noted in the figure caption.

## 3. RESULTS

### 3.1. Face-validity of individual power spectra

Our investigation assumed that an individual’s power spectrum at rest can systematically (i) differ between days, and also (ii) differ from the spectra of other individuals. We first confirmed the face-validity of these assumptions in our data.

The structured inter-individual differences during *RS1* were qualitatively evident from the mean (full) power spectrum at different channels (Figure 4A) before its reduction to the minimal description used for the classification analyses. As shown for one example individual *S*_1_, individual power spectra had a similar form across channels with a higher power in the δ and α bands and a higher overall power in the posterior and anterior channels relative to the central channels. These individual spectra also showed prominent pair-wise differences as illustrated for a few selected individuals. The diverse varieties of inter-individual differences highlight the difficulty of representing an individual’s unique properties as illustrated for individual *S*_2_. The combination of channels and frequencies (i.e., features) at which *S*_2_ and *S*_3_ showed prominent differences were not the same features at which *S*_2_ differed from *S*_5_. However, the required decision-rule to identify *S*_2_ was a <u>single</u> feature configuration capable of distinguishing *S*_2_ from *all* others while allowing *S*_2_ to be re-identified across days, despite inter-day variations.

**Figure 4:**
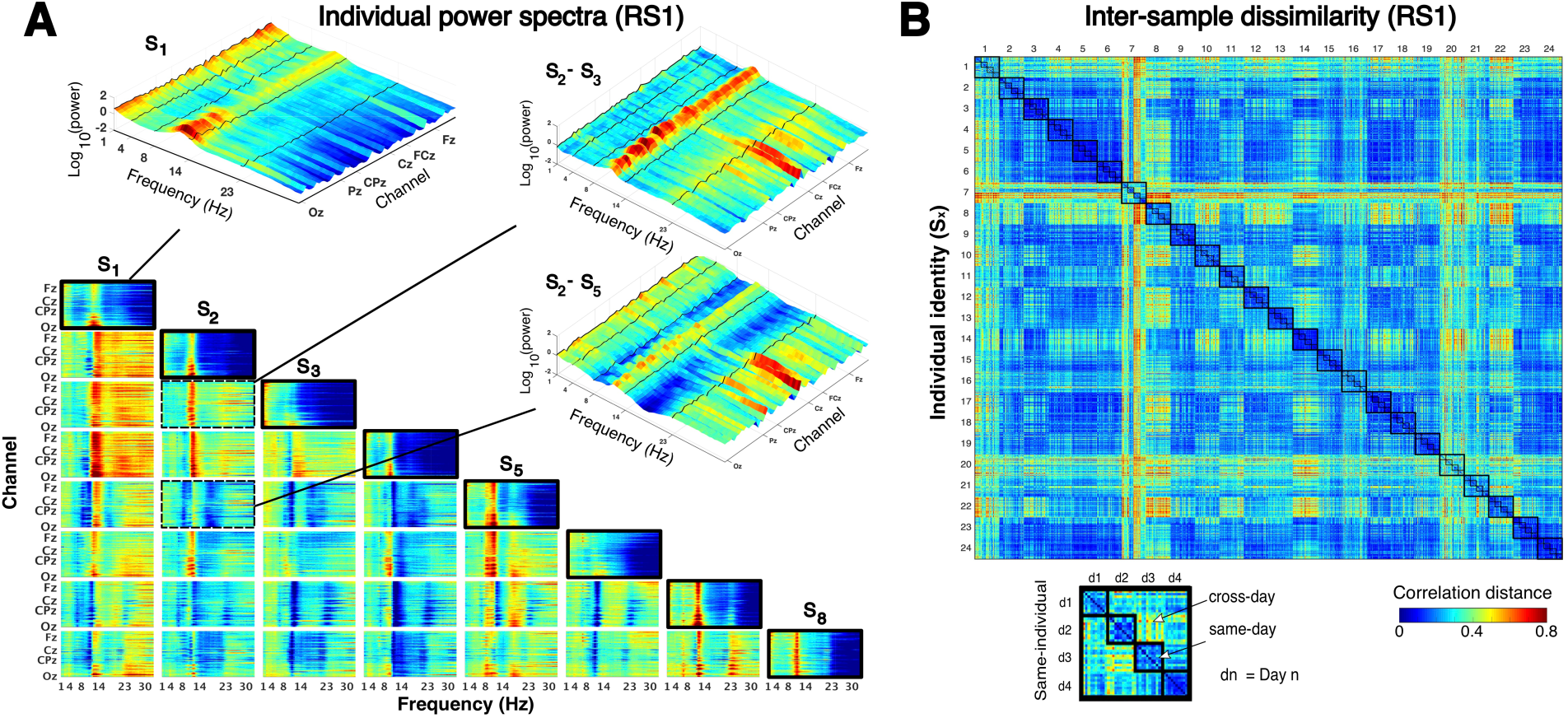
Inter-individual and inter-day differences. **(A)** Matrix showing the oscillatory (full) power spectra in *RS1* at all channels (averaged across samples and days) for 8 selected individuals (*S_i_*, diagonal, thick black boundary) and their pair-wise differences. The difference in power spectra for each pair of individuals *S_i_* and *S*_j_ (i.e., *S_i_* – *S*_j_) is shown at row *i*, column *j* of matrix. In each spectrogram, channels have a posterior-to-anterior ordering. Insets show magnified view of the power spectrum for *S*_1_ (left upper) and differences for *S*_2_ – *S*_3_ (right upper) and *S*_2_ – *S*_5_ (right lower), with frequency band boundaries marked with black lines. **(B)** Inter-sample dissimilarity matrix for *RS1* (90 samples per individual per day, each sample was defined by 305 features = 61 channels x 5 bands). The dissimilarity of two samples was defined by their correlation distance (= 1 - *r*, where *r* is the Pearson’s correlation coefficient). Large black squares on diagonal contain values from the same individual, and the four smaller squares each contain same-day values.

The systematic inter-day differences were evident from the dissimilarity between samples from all participants and all days (90 samples per participant per day) (Figure 4B). The dissimilarity between any two samples was described by their correlation distance (= 1 - *r*, where *r* is the Pearson’s correlation coefficient)(Diedrichsen and Kriegeskorte 2017; Dimsdale-Zucker and Ranganath 2019; Pani et al. 2020). For all 24 participants, the mean dissimilarity between samples from the *same day* was <u>lower</u> than between samples from different days (*cross-day*) [*t*_23_ = −6.74, p < 0.0001]. However, the dissimilarity between same-day and cross-day samples varied from person to person suggesting their possible confusability with samples from other individuals. This was the critical issue to be resolved with an appropriate decision-rule, to be identified using machine-learning.

### 3.2. Identification of individuals from RS activity within and across days

#### 3.2.1. High same-day accuracy but reduced cross-day accuracy of individual decision-rules

To identify a person from a 2s sample of RS activity with an ensemble classifier, a decision-rule was numerically estimated to represent each person’s unique RS characteristics. The decision-rules estimated for each day could identify each person (of 24) from a sample acquired on the same day (i.e., according to the scheme ^RS1^**I***_p_* → ^RS1^**I***_p_*) with a mean cross-validated (CV) accuracy of 99.98 ± 0.04% (mean ± sd) that was significantly larger than the theoretically expected accuracy for random guessing [> 50%: t_23_ = 5596.13, p < 0.00001] (Figure 5A, Table A.1). However, for longitudinal tracking, a key demand is that decision-rules from one day should identify a person from samples acquired on a different day (i.e., ^RS1^**I***_p_* → ^RS1^**I***_q_*). The same-day decision-rules identified individuals across days with a mean accuracy of 92.10% ± 6.8% that was higher than random chance [*t*_23_ = 30.14, *p* < 0.00001] but less accurate than same-day identification by ∼8% [paired *t*_23_ = 5.64, *p* = 0.00001].

**Figure 5:**
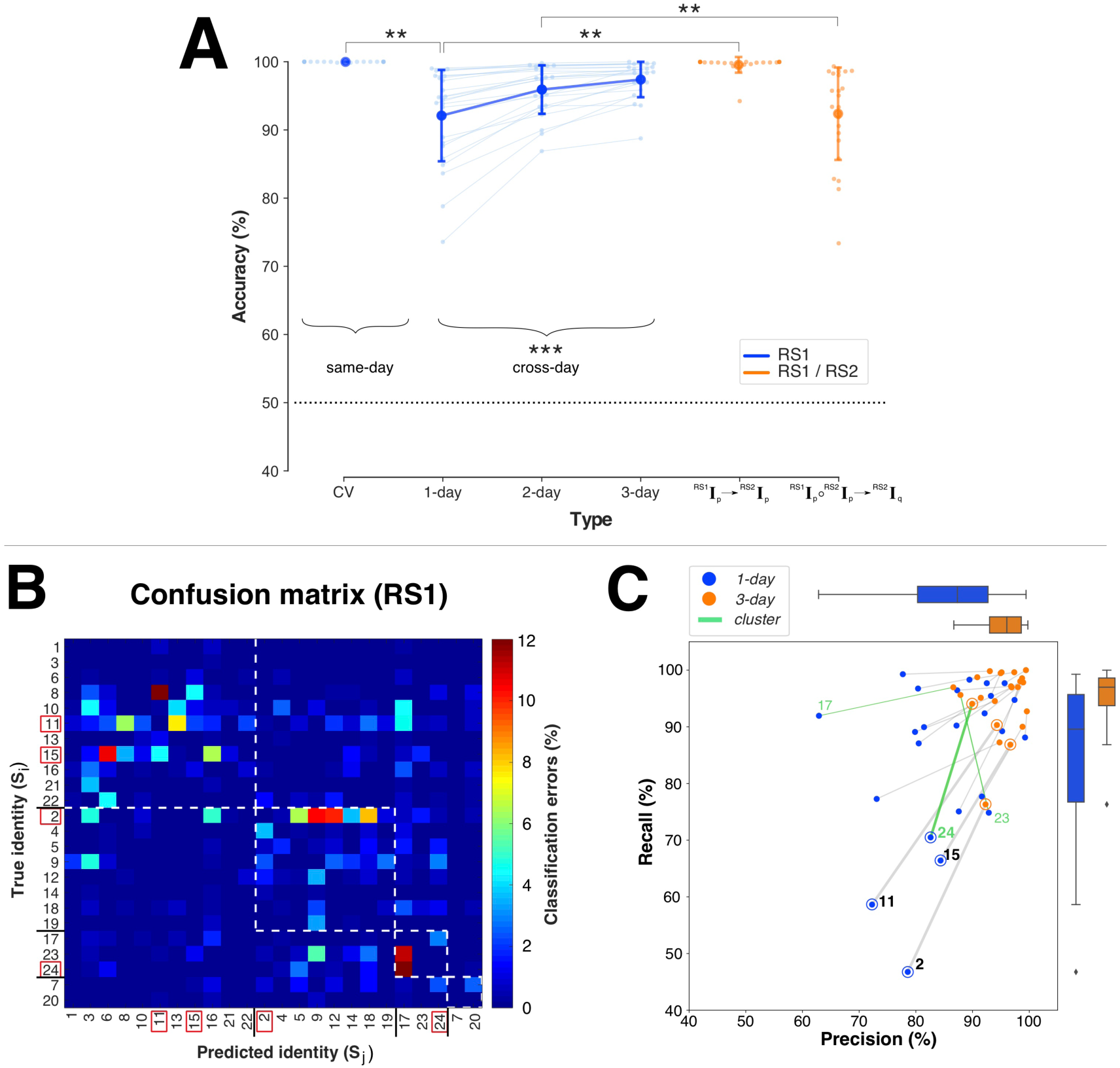
Identification accuracy at rest. **(A)** Mean identification accuracy with *RS1* (blue) on same-day (CV), across days (1-day, 2-day, 3-day), and schemes relating *RS1* and *RS2* (orange). Light colored dots/lines depict individual accuracies (*N*=24). Horizontal dotted line depicts random chance accuracy (50%). Error bars: SD (** = 0.00001 ≤ *p* < 0.001; *** = *p* < 0.00001). **(B)** Confusion matrix for cross-day (1-day) identification (only errors are shown). Dotted squares indicate clusters of individuals who are more confused with each other. Identities of individuals with the lowest cross-day accuracies are highlighted with red squares. **(C)** Changes to precision and recall scores with aggregation for the whole group (shown with boxplots) and for individuals (1-day: blue dots, 3-day: orange dots). Individuals with lowest 1-day accuracy are indicated with rings and thick gray lines. Green lines highlight *S*_24_, *S*_23_ and *S*_17_ who belong to the same cluster, shown in **(B).**

The confusion matrix (Figure 5B) of how individuals were misclassified during cross-day (1-day) identification revealed four clusters of individuals who were confused with each other. Notably, the individuals with the lowest cross-day accuracies (namely, *S*_2_*, S*_11_*, S*_15_*, S*_24_) belonged to different clusters rather than being solely confused with each other. The clustering of misclassified individuals suggested that errors in identifying an individual *S_X_* were due to a combination of (i) changes to *S*_X_’s RS-activity between days (i.e., false negatives) and (ii) changes to other individuals who were then misclassified as *S*_X_ (i.e., false positives). Nevertheless, the increased errors in individual identification illustrate the challenge of *NP+/NP-* decisions. Errors in identifying a person *S*_X_ across days seemingly imply that *S*_X_*’s* unique identifying characteristics had changed across days even though the individuals here were unlikely to have changed in their underlying neurophysiology over the 5-day testing period.

#### 3.2.2. Aggregated training increases cross-day accuracy

In numerical terms, the cross-day loss in accuracy implies that certain properties of each day’s decision-rules were of predictive relevance to same-day samples but of limited generality to other days. To discount the role of these day-specific properties in favor of day-general properties, the decision-rules were trained using samples aggregated from *multiple* days (i.e., ^RS1^**I**_p_ ∘ ^RS1^**I**_q_ … → ^RS1^**I**_s_) (Figure 5A). The mean cross-day accuracy *increased* from 92.10% ± 6.8% without aggregation (1-day) to 95.93 ± 3.63% with 2-day aggregation, with an additional increase to 97.39% ± 2.65% with 3-day aggregation [one-way ANOVA, F_2,46_ = 28.83, p < 0.00001]. Following aggregation, the cross-day accuracy was a mere ∼2% lower than the same-day accuracy. The effects of aggregated training on individual-specific identification errors are shown in Figure 5C. The decision-rules obtained with 3-day aggregation were associated with fewer false negatives (indexed by the higher recall score) especially for individuals with the lowest 1-day accuracies, i.e., *S*_2_*, S*_11_*, S*_15_ and *S*_24_. This was associated with interrelated changes in errors in individuals who belonged to the same cluster. For example, there was a prominent reduction in false positives (indexed by the higher precision score) for *S*_17_ who was in the same cluster *S*_24_ and *S*_23_ (highlighted in green). The increased accuracy with aggregation despite the true inter-day differences in RS-activity was consistent with the presence of *day-general* properties (section 2.8.4, Figure 3).

#### 3.2.3. Cross-day versus cross-measurement identification are not equivalent

We next assessed whether the above accuracy relationships across days (with and without aggregation) was related to a difference in *days* rather than simply a difference in *measurements*.

In our experimental protocol (Figure 1A), *RS2* was the second RS measurement on each day. The effects of aggregation on cross-day identification with *RS1* were successfully replicated on *RS2* without statistically detectable differences (Table A.1) [two-way ANOVA, Condition {*RS1, RS2*} x Type {1-day, 2-day, 3-day}, Type*Condition: F_2, 46_ = 0.56, p = 0.57; Type: F_2, 46_ = 31.31, p < 0.00001; Condition: F_1, 23_ = 0.38, p = 0.54]. Importantly, *RS2* validated the *day-specific* properties of the decision-rules (Figure 5A). Same-day decision-rules from *RS1* classified samples of *RS2* from the same day (^RS1^**I***_p_* → ^RS2^**I***_p_*) with a mean accuracy of 99.55 ± 1.15% that was significantly greater than the accuracy in classifying *RS1* across days (^RS1^**I***_p_* → ^RS1^**I***_q_*) (92.10 ± 6.84%) [paired t_23_ = 5.19, p = 0.00003]. Furthermore*, RS2* validated the importance of aggregating samples from different days (rather than different measurements) to reduce day-specificity. Decision-rules trained on aggregated same-day samples from *RS1* and *RS2* (^RS1^**I**_p_ ∘ ^RS2^**I**_p_ → ^RS1^**I***_r_*) had a *lower* cross-day accuracy (92.38 ± 6.92%) than decision-rules trained on aggregated *RS1* samples from two different days (^RS1^**I**_p_ ∘ ^RS1^**I**_q_ → ^RS1^**I***_r_*) (95.93 ± 3.63 %) [paired t_23_=-4.83, p = 0.00007].

In summary, the reduction in cross-day accuracy without aggregation was indicative of *inter-day* (rather than inter-measurement) variations in RS activity. Despite this inter-day variation in RS activity, the cross-day accuracy increased with aggregation and further revealed the existence of *day-general* properties in RS-activity that were unchanged across days. These properties were consistent with an activity configuration that was putatively defined by individual-specific neurophysiological constraints.

### 3.3. Information organization in resting activity enabling individual identification

The hypothesized configuration in RS-activity was suggestive of a multivariate relationship between distributed features. However, the accuracy relationships described above do not indicate whether such a distributed configuration was necessary to enable individual identification. Therefore, we evaluated the information organization required for individual identification.

#### 3.3.1. Low cross-day identification with information from only one frequency or one location

Each sample was a snapshot of RS activity described by 305 informational features (5 bands x 61 channels). To test the informational role of these different features, we evaluated whether identification comparable to the full feature-set was possible with subsets of features that were defined either by frequency band (i.e., mono-band sets) or spatial location (i.e., mono-location sets).

Each *mono-band* feature set (*B_f_)* consisted of features from one frequency band *f* at all 61 channels. For all five mono-band sets (Figure 6A, Table A.2), same-day identification had a mean accuracy greater than 95%. However, the size of the cross-day loss in accuracy was band-dependent and ranged from ∼14% for *B_α_* to nearly ∼32% for *B_δ_* [ANOVA, Type {CV, 1-day} x Band {*B_δ_, B_θ_, B_α_, B_β1_, B_β2_*}, Type*Band: F_4,92_ = 24.83, p < 0.00001; Type: F_1,23_ = 232.11, p < 0.00001; Band: F_4, 92_ = 40.30, p < 0.00001]. The divergence in cross-day losses for *B_α_* and *B_δ_* was striking as these two bands have a characteristically higher power relative to the other bands (Figure 4). Training with multi-day aggregation (Figure 6B) increased cross-day accuracy by differing amounts for each band by, for example, +10% for *B_β2_* but only +6% for *B_δ_* [ANOVA, Band {*B_δ_, B_θ_, B_α_, B_β1_, B_β2_*}x Type {*1-day, 2-day, 3-day*}, Type*Band: F_8, 184_ = 9.19, p < 0.00001; Type: F_2, 46_ = 146.02, p < 0.00001,; Band: F_4, 92_ = 43.13, p < 0.00001]. However, even with 3-day aggregation, the residual difference between cross-day and same-day accuracy (minimum: ∼7% for *B_α_*, maximum: ∼26% for *B_δ_*) was larger than the ∼2% difference with the full feature-set.

**Figure 6:**
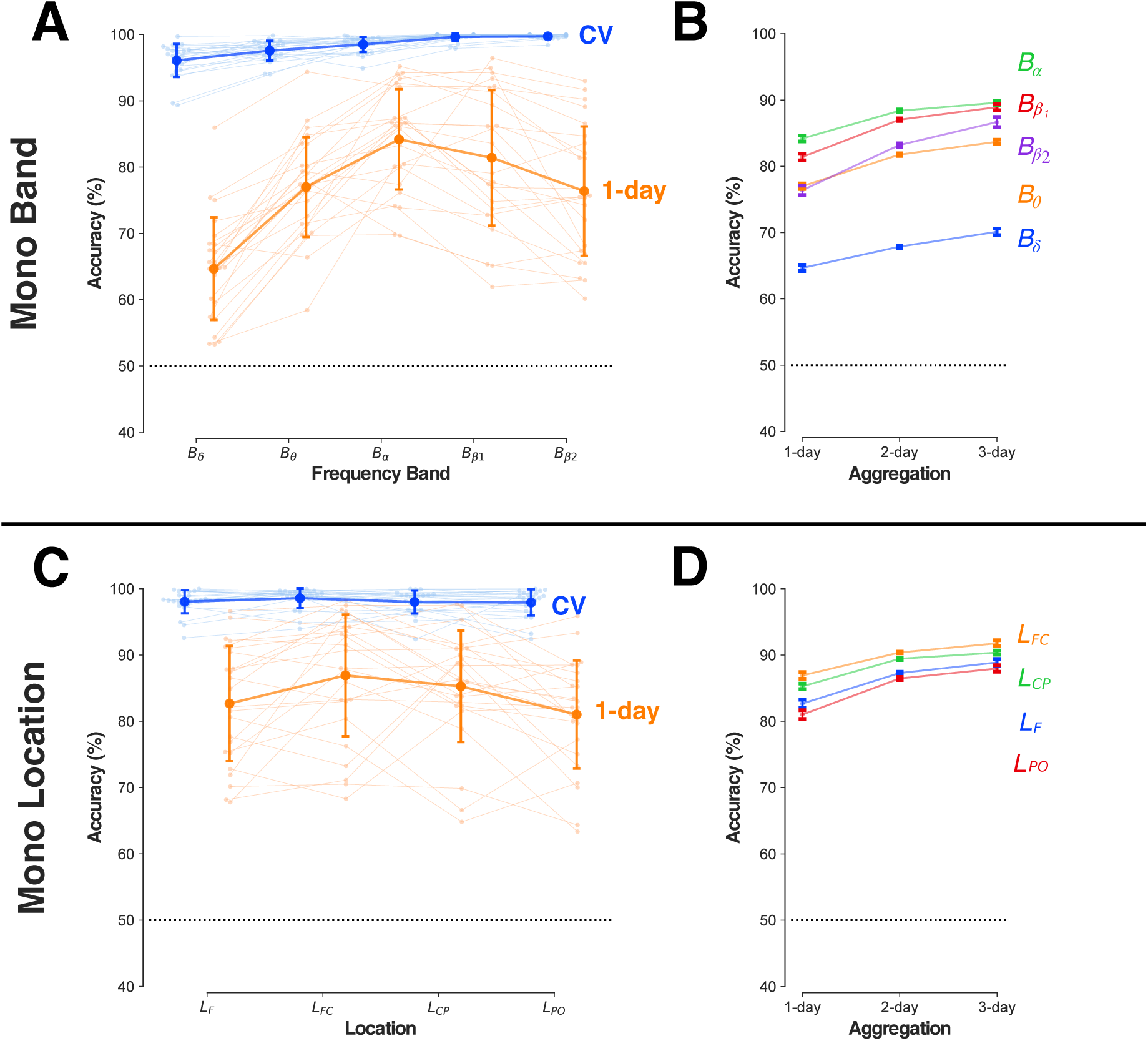
Identification at rest with mono-band/location feature subsets. **(A)** Mean identification accuracy for *RS1* with mono-band feature sets of increasing frequency (x-axis) on the same-day (blue, CV) and across-days (orange, 1-day). Light-colored dots/lines depict individual accuracies (*N*=24). Error bars: SD. **(B)** Change in cross-day identification with increasing aggregation (x-axis) for different mono-band feature subsets (colored lines). Error bars: Within-subject s.e.m. (O’Brien and Cousineau 2014). **(C)** Mean identification accuracy for mono-location feature sets (x-axis, from anterior to posterior) with graphical representation and error bars as in panel **A**. **(D)** Change in cross-day identification with increasing aggregation (x-axis) for different mono-location feature subsets (colored lines). Error bars: Within-subject s.e.m. Horizontal dotted lines depicts the random chance accuracy (50%) in all panels.

Each *mono-location* feature set (*L_z_*) consisted of 50 features (5 bands x 10 channels) in the spatial zone *z* (Figure 2A). The mean same-day accuracy was greater than 95% for all mono-location feature sets (Figure 6C, Table A.2). However, the mean cross-day (1-day) accuracy showed reductions of ∼12%-16% for all locations [ANOVA, Type {CV, 1-day} x Location {*L*_F_, *L*_FC_, *L*_CP_, *L*_PO_}, Type*Location: F_3,69_ = 3.77, p = 0.015; Type: F_1,23_ = 108.91, p < 0.00001,; Location: F_3, 69_ = 5.45, p = 0.0020]. The mean cross-day accuracy for the fronto-central (*L*_FC_) and centro-parietal (*L*_CP_) sets were marginally higher than for the parieto-occipital (*L*_PO_) and frontal (*L*_F_) sets. This zonal accuracy difference was notable as the mean power for all bands was typically higher over the posterior and anterior channels than the centrally located channels (Figure 4A). Aggregation increased cross-day accuracy by ∼6% for all four location sets (Figure 6D) [ANOVA, Location: {*L*_F_, *L*_FC_, *L*_CP_, *L*_PO_} x Type {*1-day, 2-day, 3-day*} [Type*Location: F_6, 138_ = 2.07, p = 0.06; Type: F_2, 46_ = 115.38, p < 0.00001; Location: F_3, 69_ = 4.79, p = 0.0043]. Nevertheless, the residual ∼7%-10% loss in cross-day accuracy was larger than with the full feature-set.

In summary, all the mono-band and mono-location sets contained sufficient information to enable same-day identification with nearly error-free accuracy. However, this information had a low day-generality. Even with aggregation, these feature sets had a much lower cross-day accuracy than the full feature-set that combined these feature sets together. This is notable with regard to machine learning algorithms. Generalization accuracy can reduce with an increase in the number of features (the so called Hughes effect (Campenhout 1978; Sima and Dougherty 2008)). However, here, a feature set of 305 features showed greater cross-day generalization than small feature-sets of 50/60 features that have comparable same-day cross-validated accuracy. This divergence suggests that the higher cross-day robustness with the full feature-set involves a role for multivariate relationships between different frequency bands (i.e., unlike the mono-band subsets) at spatially distributed channels (i.e., unlike the mono-location subsets). To assess how this multi-feature configuration was organized, we evaluated the pattern of weights associated with the different features of the full feature-set.

#### 3.3.2. Concentration of high-consistency features at fronto-central and occipital zones

Each individual’s linear decision-rule was defined by the configuration of weights assigned to the different features, where weights with a larger magnitude (irrespective of sign) have a larger role in the classification decision even if in an indirect manner (Haufe et al. 2014; Schrouff and Mourao-Miranda 2018). However, individual-specific weight configurations might differ from each other in an idiosyncratic manner with little consistency between individuals since, for example, a high-weighted feature in *S*_X_’s decision-rule might be of limited relevance to individual *S*_Y_’s decision-rule.

Figure 7 shows the topographic distribution of high-consistency features in the full feature-set after normalization for power differences (see Suppl. Figure 1 for high-consistency non-normalized (raw) weights). At corrected thresholds (see *t-*values in Figure 7, lower panels), the features associated with all frequency bands except the *δ* band contained at least one high-consistency feature. Rather than having an idiosyncratic organization, the high-consistency features were concentrated at distinctive zones in each frequency band.

**Figure 7:**
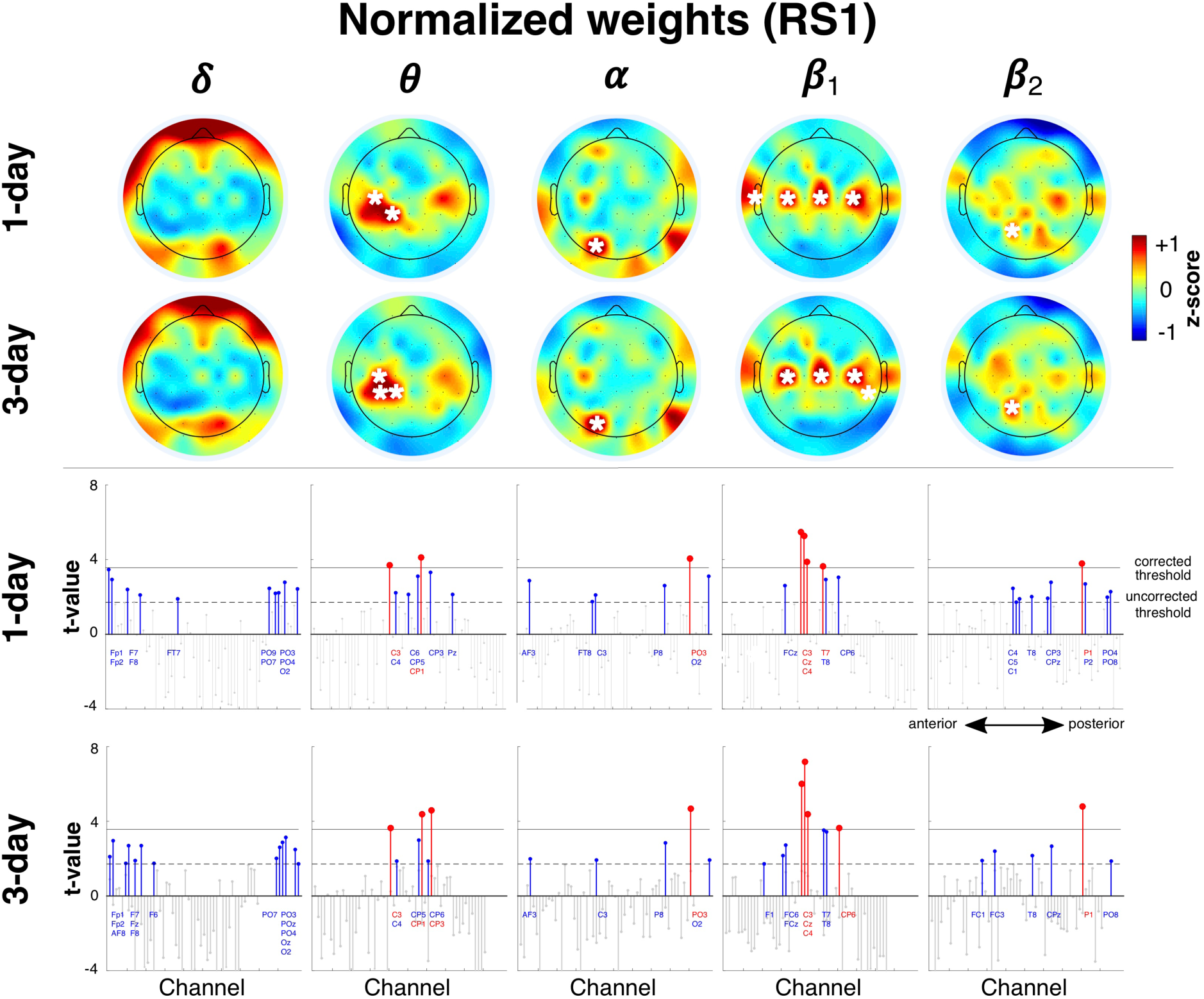
High-consistency features. Spatial distribution of high-consistency normalized weights for frequency bands of full feature set (z-scored per band) and their changes with aggregation (1-day, 3-day). Mean weights in each scalp map that were significantly greater than zero are indicated with a white asterisk (*p* < 0.05/61). Lower two rows show *t*-values for the features corresponding to the upper rows. Channels have an anterior-to-posterior ordering (x-axis). Red stems indicate channels with t-values higher than the corrected threshold (*p* < 0.05/61, horizontal black line) while blue stems show channels that only pass uncorrected thresholds (*p* < 0.05, dotted horizontal line). Colored channel labels are grouped from top-to-bottom for visibility and correspond to stems from left to right.

In *B_θ_*, there was a concentration of high consistency features at CP1 and C3, with the addition of CP3 with aggregation. There was a similar, although weaker, concentration of consistent features at corresponding channels over the right hemisphere. Showing a similar spatial organization, the high-consistency features in *B_β1_* showed a striking bilaterally symmetric configuration along the transverse midline at channels C3, Cz and C4 with an aggregation-modulated role for CP6 and T7 (and possibly T8). This similarity in organization was notable since the frequency ranges of the *θ* band (4-7.5 Hz) and β_1_ (14-22.5 Hz) were not contiguous and were separated by the *α* band.

Unlike this central concentration of features in *B_β1_* and *B_θ_*, the features in *B_α_* contained a single, strongly consistent feature in the occipital zone at PO3. At uncorrected thresholds, there were other distributed features across the scalp that were weakly consistent for both 1-day and 3-day identification, namely, at AF3, C3, P8 and O2. Similarly, the features of the high-frequency *β_2_* band (i.e., *B_β2_*) only had a single consistent feature at P1 with a diffuse distribution of consistent features at uncorrected thresholds.

In general, the distribution of high-consistency features was by itself not a simple indicator of their contribution to cross-day accuracy. For example, the relative number of high-valued weights in the different bands and spatial locations had a low correspondence to relative accuracy of cross-day identification based solely on the mono-band/location subsets (see Supplementary Figure 2). Nevertheless, the organized distribution of high-consistency features at channels over the sensorimotor cortex and the occipital cortex was prima facie support for an individual-specific configuration with a basis in neurophysiological constraints. The relevance of the high-consistency zones was of particular interest to the relationship of *RS1* to the non-rest task states where the power over the sensorimotor and occipital zones was expected to differ from *RS1*.

### 3.4. Relationship of rest to non-rest states

The behavioral demands during *TapMov* and *SeqMov* were designed to modulate the cognitive states during the *TapWait* and *SeqWait* periods and produce neural activity deviations from *RS1* in the absence of behavioral differences. Furthermore, the *Tap* and *Sequence* tasks were designed to elicit neural states that varied between days for *Sequence* (low cross-day similarity) but remained constant for *Tap* (high cross-day similarity). We sought to first explicitly verify that such deviations from *RS1* were indeed present. Note that all analyses of *Tap* and *Seq* states were performed in a subgroup of *N=*18 participants (see section 2.6).

#### 3.4.1. Neural activity during Tap and Sequence verifiably deviate from RS1

The inter-day changes in behavior during the *TapMov* and *SeqMov* periods were consistent with the experimental assumptions (Figure 8A). During *TapMov*, the mean number of button presses per trial (∼12-13) remained effectively constant across days [one-way ANOVA, F_4, 68_ = 0.55, p = 0.70]. In contrast, during *SeqMov*, the mean number of button-presses increased from ∼8 on the first day to ∼13 on the fifth day [one-way ANOVA, F_4, 68_ = 21.36, p < 0.00001]. This inter-day change in motor performance in *SeqMov* was systematically different from *TapMov* as confirmed by the statistically significant interaction in an ANOVA with factors Condition {*TapMov*, *SeqMov*} x Days {D1,..,D5} [Condition*Days: F_4, 68_ = 12.38, p < 0.00001; Condition: F_1, 17_ = 3.13, p = 0.095; Days: F_4, 68_ = 10.71, p < 0.00001].

**Figure 8:**
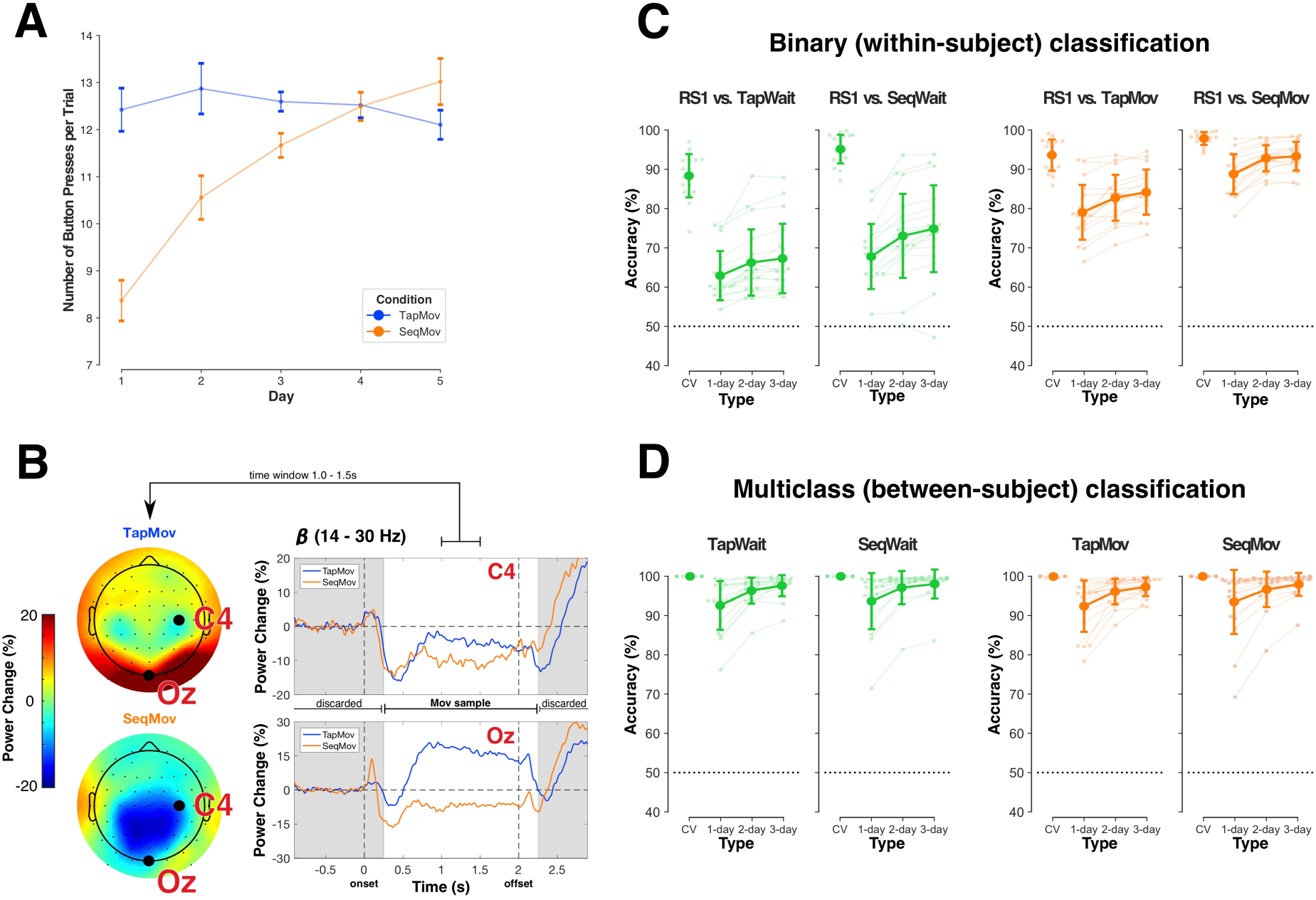
Difference between *RS1* and task-states. **(A)** Change in behavior indexed by mean number of button-presses across days (x-axis) during *TapMov* (blue) and *SeqMov* (orange). Error bars: Within-subject s.e.m. **(B)** Movement-related power dynamics in the *β* band (14-30 Hz) in *TapMov* (blue) and *SeqMov* (orange) at channels C4 (upper right) and Oz (lower right) averaged across participants and days. Intervals marked in gray were discarded from the *TapWait* and *SeqWait* samples used for classification to avoid movement-related carry over effects into the waiting periods. Scalp plots (left panel) show the mean power distribution over the period [+1s, +1.5s] following onset of the movement cue. **(C)** Same-day/cross-day accuracy in distinguishing *RS1* vs pseudo-rest states (green) and *RS1* vs movement states (orange) using within-subject binary classifiers. Cross-day differences to *RS1* were lowest for *TapWait* (far left) and highest for *SeqMov* (far right). Error bars: SD. (**D**) Person identification accuracy (multiclass) when the training/test sets were from the same task state (green: pseudo-rest states, orange: movement states). Error bars: SD.

The neural state during the movement-period (*TapMov, SeqMov)* showed typically expected dynamic states (Figure 8B). Changes in the mean *β* power at channel C4 (contralateral to the moved fingers) were in line with the Event-Related De-synchronization/Synchronization (ERD/ERS) phenomenon for repetitive movements (Pfurtscheller and Lopes da Silva 1999; Cassim et al. 2000; Alegre et al. 2004; Erbil and Ungan 2007), namely, a power reduction at the onset of movement execution (i.e., ERD) with an increase after the termination of all movements (i.e., ERS). Furthermore, the *β* power changes at Oz showed a task-dependent neural response consistent with differing visual stimulation, that is, an increase for *TapMov* (blank screen) but a decrease for *SeqMov* (image depicting the sequence). These movement-vs-wait differences were validated in the samples used for classification. A within-subject binary classification of *TapWait* vs *TapMov* had a mean cross-validated accuracy of 85.91 ± 7.23% [> 50%: t_17_ = 21.06, p < 0.00001]; and *SeqWait* vs *SeqMov* had a mean CV accuracy of 94.58 ± 3.20% [> 50%: t_17_ = 59.02, p < 0.00001].

The critical verification for our study was the relationship between *RS1* and the pseudo-rest states (*TapWait*, *SeqWait*). Samples from *TapWait* and *SeqWait* were distinguishable from *RS1* on the same day with high cross-validated accuracy (*RS1* vs *TapWait*: 88.28 ± 5.70%; *RS1* vs *SeqWait*: 95.12 ± 3.74%) (Figure 8C left panels, Table A.3). However, the cross-day accuracy (without aggregation) for both *RS1* vs *TapWait* (62.91 ± 6.44%) and *RS1* vs *SeqWait* (67.79 ± 8.53%) was substantially lower than the same-day accuracy by more than ∼25%. Nevertheless, the cross-day accuracy for *RS1* vs *SeqWait* was marginally higher than for *RS1* vs *TapWait* with increasing aggregation [ANOVA: Condition {*RS1* vs *TapWait*, *RS1* vs *SeqWait*} x Type {1-day, 2-day, 3-day}, Condition*Type: F_2, 34_ = 6.22, p = 0.005; Condition: F_1, 17_ = 8.37, p = 0.01009; Type: F_2, 34_ = 38.89, p < 0.00001].

*TapMov* and *SeqMov* were also distinguishable from *RS1* on the same-day with high (cross-validated) accuracy (*RS1* vs *TapMov:* 93.56 ± 4.12%; *RS1* vs *SeqMov*: 97.81 ± 1.76%) (Figure 8C, right panel, Table A.3). Similar to the wait periods, the cross-day accuracy for *RS1* vs *SeqMov* was higher than for *RS1* vs *TapMov* across aggregation levels [ANOVA: Condition {RS vs *TapMov*, RS1 vs *SeqMov*} x Type {1-day, 2-day, 3-day}, Condition*Type: F_2, 34_ = 0.61, p = 0.55; Condition: F_1, 17_ = 30.91, p = 0.00003; Type: F_2, 34_ = 69.47, p < 0.00001].

The above findings verified the neural activity differences in the task-states in *Tap* and *Sequence* to each other and to *RS1*. Crucially, the structure of the same-day differences had a low cross-day generality.

#### 3.4.2. Robust identification of individuals from Tap and Sequence activity within and across days

The above differences between task-states and *RS1* raised the issue of whether the task-related functional states also disrupt the information that enables individual identification with *RS1.* To assess this possibility, we evaluated whether the different *Tap* and *Sequence* task-states contained sufficient information for person identification in a same-task classification scheme (i.e., with the scheme ^X^**I**_p_ → ^X^**I**_q_ for task *X*) (Figure 8D).

The same-day accuracy for both *TapWait* and *SeqWait* was ∼99% (Figure 8D left panels, Table A.1). The mean cross-day accuracy (without aggregation) for *TapWait* (92.58 ± 6.39%) was lower than its corresponding same-day accuracy by only ∼7% [t_17_ = 4.92, p = 0.00013]. Similarly, for *SeqWait*, the mean cross-day (1-day) (93.67 ± 7.35%) accuracy was lower than the same-day accuracy by ∼6% [t_17_ = 3.65, p = 0.00197]. Furthermore, the effect of aggregation on mean cross-day accuracy for *TapWait* and for *SeqWait* were statistically indistinguishable [ANOVA: Condition {*TapWait*, *SeqWait*} x Type {1-day, 2-day, 3-day} [Condition*Type: F_2, 34_ = 0.88, p = 0.42; Condition: F_1,17_ = 1.35, p = 0.26; Type: F_2, 34_ = 21.30, p < 0.00001].

Despite the deviations of *TapMov* and *SeqMov* along both the behavioral and cognitive dimensions of rest and their differences with each other, the accuracies of individual identification across days for *TapMov* and *SeqMov* were greater than 90% for all levels of aggregation and were not statistically distinguishable from each other (Table A.1, Figure 8D right panels) [ANOVA: Condition {*TapMov*, *SeqMov*} x Type {1-day, 2-day, 3-day} [Condition*Type: F_2, 34_ = 0.86, p = 0.43; Condition: F_1, 17_ = 1.26, p = 0.28; Type: F_2, 34_ = 14.50, p = 0.00003].

Thus, individual identification was robustly possible in the task states despite their differences to *RS1*. Furthermore, the identification accuracy was similar between the *Tap* and *Seq* states despite their functional differences. Two further lines of evidence supported the possibility that these similarities were based on common task-independent properties. The spatial distribution of high-consistency features for these states (Figure 9A, Suppl. Figure 3) exhibited a striking qualitative similarity to each other as well as to the corresponding distribution for *RS1* (Figure 7). Additionally, the individual cross-day (1-day) accuracy in these task states showed a striking correlation to the corresponding cross-day accuracy in *RS1* (Figure 9B)[threshold: *p* < 0.05/4; *TapWait*: *r*(17)=0.882, *p* < 0.00001; *SeqWait*: *r*(17)=0.635, *p* = 0.00466; *TapMov*: *r*(17)=0.75, *p* = 0.00034; *SeqMov*: *r*(17)=0.653, *p* = 0.00329]. Thus, the inter-individual relationships revealed by the errors in cross-day classification during *RS1* (Figure 5B) seemingly extended to these non-rest states as well. We next turned to a formal assessment of this cross-task relationship.

**Figure 9:**
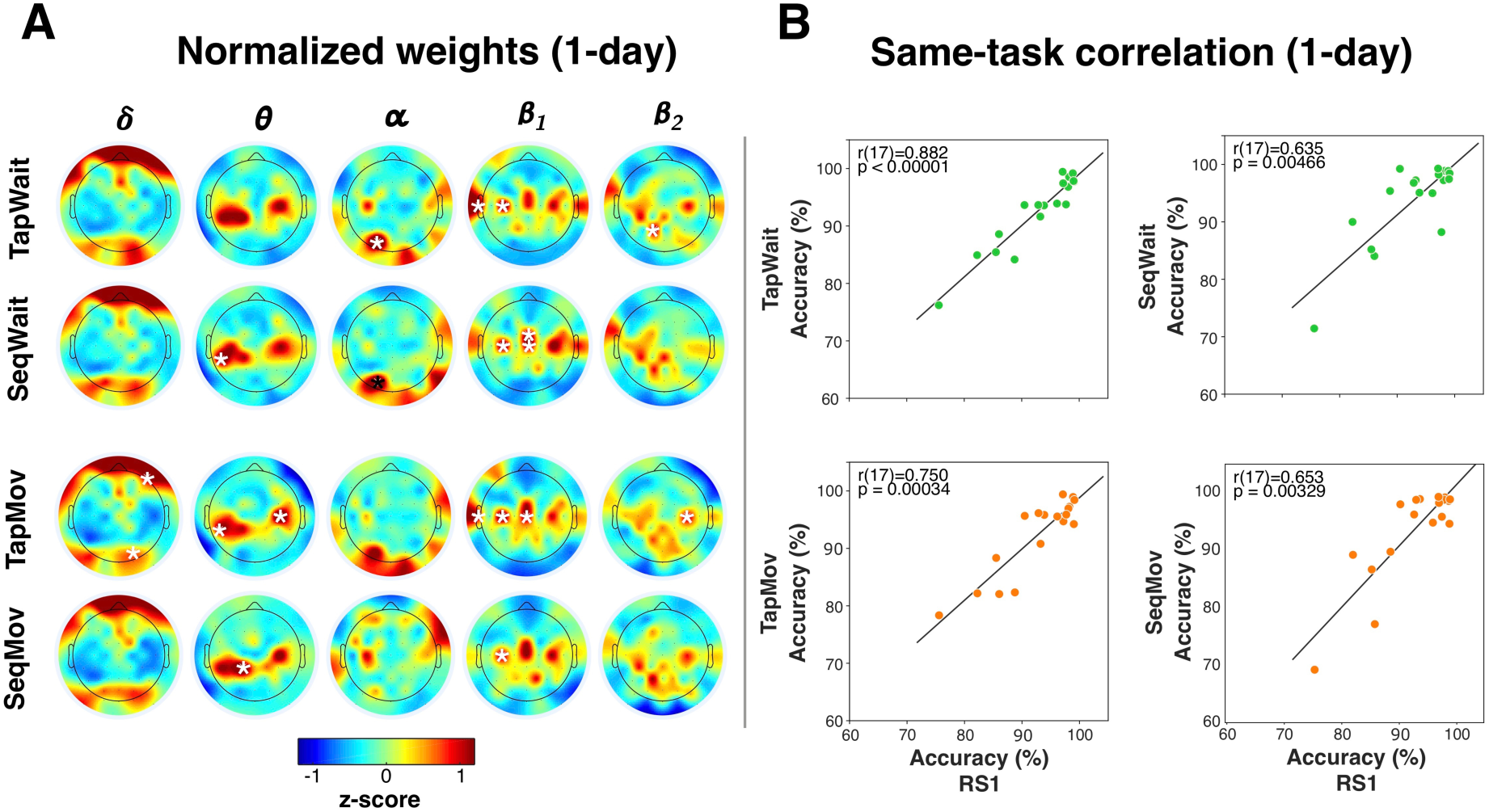
Inter-task relationships. **(A)** Spatial distribution of high-consistency features in different task-states (absolute, z-scored) for 1-day decision-rules (without aggregation). Weights in each scalp map that were significantly greater than zero are indicated with a white asterisk (*p* < 0.05/61, see Supplementary Figure 3). Each frequency band (column) had a characteristic spatial distribution of high weighted channels that was qualitatively similar across task-states and also to *RS1* (Figure 7). **(B)** Scatter plots of cross-day (1-day) identification accuracy in *RS1* to the corresponding same-task accuracy in the pseudo-rest states (upper row) and movement states (lower row). Each dot represents one individual. Correlations were assessed with Spearman’s rank order correlation (threshold: *p* < 0.05/4).

### 3.5. Generalization of rest-based decision to cross-task individual identification

If person identification with *RS1* was based on a neural configuration related to an individual’s neurophysiological state then identification should be possible despite cognitive state variations. Therefore, decision-rules trained on *RS1* should be capable of accurate person identification with samples acquired from the pseudo-rest states (*TapWait* and *SeqWait*) and the movement states (*TapMov* and *SeqMov*).

#### 3.5.1. Robust cross-task identification with RS1 with full feature-set

We used the cross-task scheme ^RS1^**I**_p_ → ^X^**I**_q_ to test the invariance of *RS1-*based identification to inter-day cognitive state variations (i.e., task states *X*) (Figure 10A, Table A.4). Increasing deviations from *RS1* solely due to cognitive state differences (*X = {RS1, TapWait, SeqWait}*) did not produce comparable, statistically distinguishable reductions in mean identification accuracy (*RS1*: 92.79 ± 6.76%, *TapWait*: 91.90 ± 6.46%; *SeqWait*: 90.81 ± 7.09%) [one-way ANOVA, F_2, 34_ = 2.06, p = 0.14]. However, increasing deviations from *RS1* due to cognitive <u>and</u> behavioral state differences (*X = {RS1, TapMov, SeqMov}*) produced significant reductions in identification accuracy most notably for *SeqMov* (*TapMov*: 88.79 ± 7.57%; *SeqMov*: 83.85 ± 10.35%)[one-way ANOVA, F_2, 34_ = 14.07, p = 0.00004].

**Figure 10:**
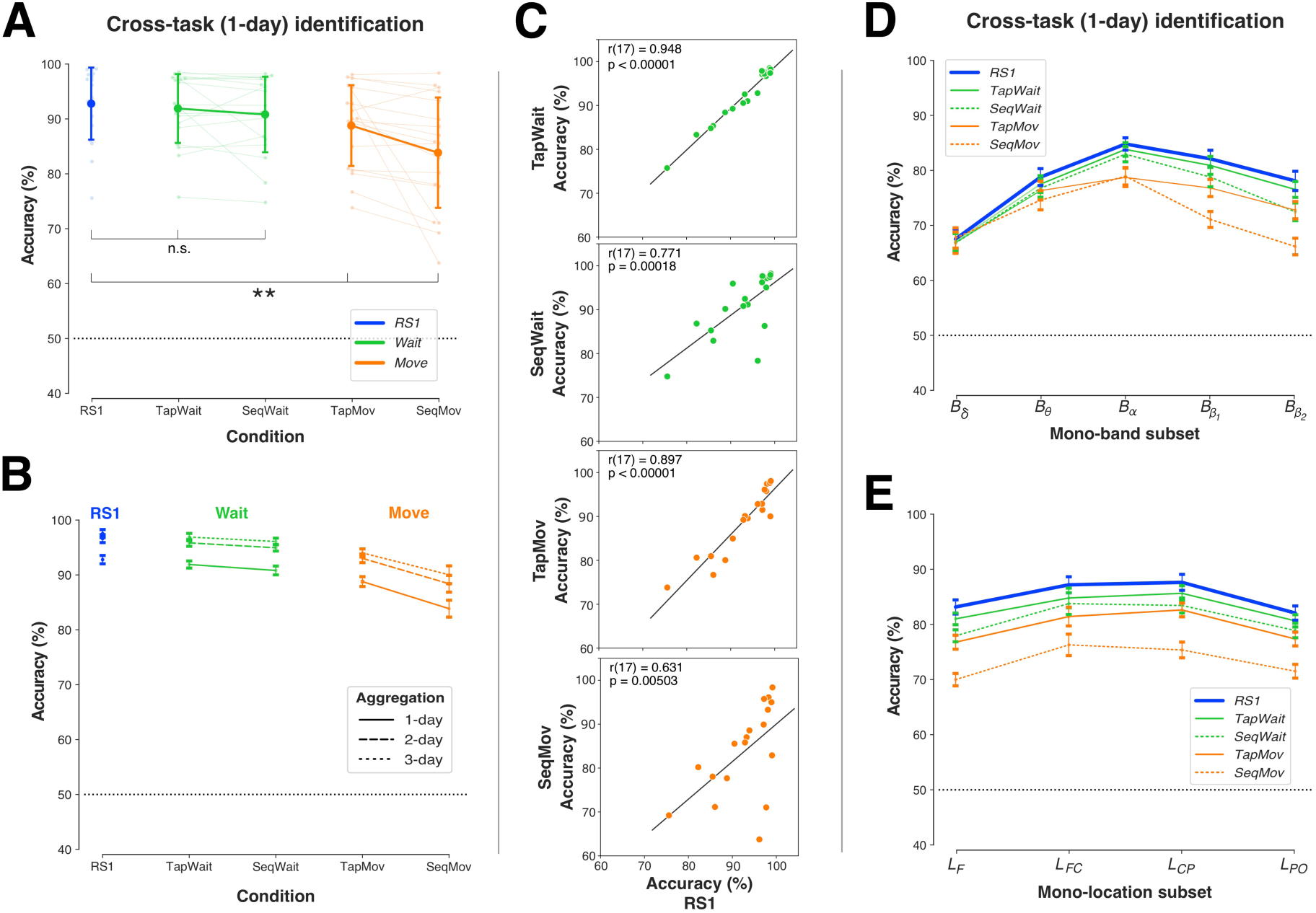
Cross-task identification with RS1. **(A)** Mean accuracy of decision-rules trained on *RS1* (1-day) and tested across days on *RS1* (blue), pseudo-rest states (green) and movement states (orange). Light colored dots/lines indicate individual accuracies. Error bars: SD. Accuracy differences between *RS1* and pseudo-rest states were not statistically significant (n.s.), but were between *RS1* and movement states (** = 0.00001 ≤ *p* < 0.001). (**B**) Decision-rules trained on *RS1* with different levels with aggregation (dotted lines) increased cross-day accuracy for all task-states. Error bars: Within-subject s.e.m. (**C**) Scatter plots of cross-day (1-day) accuracy in *RS1* to the corresponding cross-task accuracy in all non-rest tasks. Each dot represents one individual. Correlations were assessed with Spearman’s rank order correlation (threshold: *p* < 0.05/4). **(D)** Cross-task/day accuracy of *RS1* with mono-band subsets. Deviations from cross-day accuracy for *RS1* were larger for the movement states (orange) than the pseudo-rest states (green) and deviations increased with the frequency (lowest for *B*_δ_, highest for *B*_β2_). Error bars: Within-subject s.e.m. **(E)** Cross-task/day accuracy of *RS1* with mono-location subsets. Deviations from *RS1* were larger for movement states (orange) than pseudo-rest states (green). Error bars: Within-subject s.e.m.

To disentangle the role of cross-task from cross-day effects, we compared cross-task (^RS1^**I**_p_→^X^**I**_q_) and same-task identification (^X^**I**_p_ → ^X^**I**_q_) across days (Table A.1, 4). For the pseudo-rest states (*X = {TapWait, SeqWait}*), cross-task accuracy with *RS1* decision-rules produced a small but statistically significant reduction relative to same-task identification [ANOVA, Train {*RS1*, *Same*} x Condition {*TapWait*, *SeqWait*}, Train*Condition: F_1,17_ = 4.14, p = 0.06; Train: F_1,17_ = 10.02, p = 0.00566; Condition: F_1, 17_ = 0.00001, p = 1.00]. The cross-task accuracy reduction was significantly larger for the movement-states (*X = {TapMov, SeqMov}*) with a larger loss for *SeqMov* [ANOVA, Train{*RS*,*Same*} x Condition {*TapMov*, *SeqMov*}, Train*Condition: F_1,17_ = 9.15, p = 0.00764; Train: F_1,17_ = 43.94, p < 0.00001; Condition: F_1, 17_ = 2.51, p = 0.13].

To disentangle the role of day-specificity in ^RS1^**I**_p_ → ^X^**I**_q_, we used multi-day aggregation (^RS1^**I**_p_ ∘ ^RS1^**I**_q_ … → ^X^**I**_r_). Although aggregation reduced day-specificity with *RS1* (Figure 5), it could nevertheless increase specificity to the properties of *RS1*. If so, aggregation might lower the accuracy of cross-task identification. Contrary to this possibility, aggregation *increased* cross-task accuracy to the pseudo-rest states (*TapWait*, *SeqWait*) in a comparable manner to same-task accuracy (Figure 10B) [ANOVA: Condition {*RS1, TapWait, SeqWait*} x Type {1-day, 2-day, 3-day} [Condition*Type: F_4, 68_ = 0.52, p = 0.72; Condition: F_2, 34_ = 2.44, p = 0.10; Type: F_2, 34_ = 21.63, p < 0.00001]. This was particularly striking because aggregation (i.e., related to day-specificity) produced a relatively larger increase in cross-task accuracy than a change in task-specificity. Following aggregation, the mean residual cross-task/day accuracy loss relative to same-task/day identification with *RS1* was only ∼3%. Aggregation also *increased* cross-task accuracy to the movement states (*TapMov, SeqMov*) [ANOVA: Condition {*RS1, TapMov*, *SeqMov*} x Type {1-day, 2-day, 3-day} [Condition*Type: F_4, 68_ = 1.35, p = 0.26; Condition: F_2, 34_ = 13.04, p = 0.00006; Type: F_2, 34_ = 29.33, p < 0.00001]. Following aggregation, the mean residual cross-task/day difference was less than ∼10% for the movement states.

Similar to the same-task correlations described above (section 3.4.2, Figure 9B), the individual *cross-task* (1-day) accuracy in each of these task states showed a statistical significant correlation to the corresponding cross-day accuracy in *RS1* (Figure 10C)[threshold: *p* < 0.05/4; *TapWait*: *r*(17)=0.948, *p* < 0.00001; *SeqWait*: *r*(17)=0.771, *p* = 0.00018; *TapMov*: *r*(17)=0.897, *p*<0.00001; *SeqMov*: *r*(17)=0.631, *p* = 0.00503]. The correlation coefficients were particularly high for both *Tap* states *(TapWait* and *TapMov*) as compared to the *Seq* states (*SeqWait* and *SeqMov),* Furthermore, the scatter plots suggested that the relatively lower cross-task/day accuracy for *SeqMov* was driven by the low generalization of a few individuals.

In summary, decision-rules trained on *RS1* on a single day could identify individuals from samples from states that verifiably differed from *RS1* to differing extents. Importantly, aggregated training solely on *RS1* lead to *increases* in identification accuracy on samples from these non-rest task states.

#### 3.5.2. Low cross-task identification with feature subsets

The full-feature set has a crucial role in limiting the cross-day loss in accuracy in *RS1* (Figure 6). Appling the cross-task scheme ^RS1^**I**_p_ → ^X^**I**_q_ to the mono-band (Figure 10D) and mono-location (Figure 10E) feature sets provided further evidence of the importance of the full feature-set to enable robust cross-task identification.

For the mono-band feature sets (Figure 10D), increasing deviations from *RS1* in cognitive state (*X = {RS1, TapWait, SeqWait}*) lead to state-related accuracy reductions that were also larger for the higher frequency bands [ANOVA: Condition {*RS1, TapWait, SeqWait*} x Band {*B_δ_, B_θ_, B_α_, B_β1_, B_β2_*} [Condition*Band: F_8, 136_ = 2.60, p = 0.01136; Condition: F_2, 34_ = 6.44, p = 0.00426; Band: F_4, 68_ = 18.36, p < 0.00001]. In a similar manner, increasing deviations from *RS1* for the movement states (*X = {RS1, TapMov, SeqMov}*) produced state-related accuracy reductions that were greater for *SeqMov* than for *TapMov* particularly at the higher frequencies [ANOVA: Condition {*RS1, TapMov, SeqMov*} x Band {*B_δ_, B_θ_, B_α_, B_β1_, B_β2_*} [Condition*Band: F_8, 136_ = 8.95, p < 0.00001; Condition: F_2, 34_ = 20.59, p < 0.00001; Band: F_4, 68_ = 12.16, p < 0.00001]. These task-linked accuracy reductions were notably absent at *B_δ_*.

The pattern of cross-task accuracy deviation from *RS1* took a different form for the mono-location feature sets (Figure 10E). For the pseudo-rest states (*X = {RS1, TapWait, SeqWait}*), increasing deviations from *RS1* lead to increasing accuracy reductions (largest for *SeqWait*) that were relatively uniform at all the locations [ANOVA: Condition {*RS1, TapWait, SeqWait*} x Location {*L*_F_, *L*_FC_, *L*_CP_, *L*_PO_} [Condition*Location: F_6, 102_ = 0.98, p = 0.45; Condition: F_2, 34_ = 8.99, p = 0.00073; Location: F_3, 51_ = 4.34, p = 0.00849]. This pattern of reduction was similar for the movement states (*X = {RS1, TapMov, SeqMov}*), where deviations from *RS1* lead to accuracy reductions that were largest for *SeqMov* and relatively uniform at all locations [ANOVA: Condition {*RS1, TapMov, SeqMov*} x Location {*L*_F_, *L*_FC_, *L*_CP_, *L*_PO_} [Condition*Location: F_6, 102_ = 0.99, p = 0.44; Condition: F_2, 34_ = 32.92, p < 0.00001; Location: F_3, 51_ = 4.95, p = 0.00432].

Thus, the large accuracy reductions with band/location-defined feature subsets confirmed that the full feature-set was crucial to high cross-task identification accuracy. Taken together, the cross-task/cross-day robustness of person identification with the full feature-set was consistent with the hypothesized properties of a configuration constrained by individual neurophysiology.

## 4. DISCUSSION

The central motivation for the current study was whether RS-activity could support a critical demand for individualized longitudinal tracking, namely, decoding the origin of inter-day RS differences (i.e., *NP+* or *NP-*) from the relationship between the resting state activity patterns. A major obstacle to *NP+/NP-* decoding was the ill-defined rest task itself and its potential to confound the interpretation of RS-activity *differences*. To evaluate a *commonality-*based alternative, we hypothesized that the existence of an activity configuration defined by neurophysiological constraints would afford an escape from the confounding effects of the rest task. Our findings support the existence of such a configuration in the longitudinal characteristics of the EEG oscillatory power spectrum at rest. Formulated in terms of individual identification, inter-day differences in individual RS-activity were classified with high accuracy across a diverse range of confounding inter-day differences, with day-generality confirmed using aggregation. Consistent with a configuration based in whole-brain neurophysiology, accurate identification was higher with a full feature-set that enabled the integration of information from multiple frequency bands at channels distributed across the scalp.

### 4.1. Empirical simulations of cognitive and neurophysiological variation

A methodological novelty here was our use of empirical “simulations”. Although ad hoc, they provided a means to obtain verifiable instances of cognitive state variation and neurophysiological change relative to RS.

As previous studies have demonstrated (Duncan and Northoff 2013; Kawagoe et al. 2018), the potential for arbitrary cognitive state variation during the rest task is related to experimental context and instructions. However, beyond the assumption that participants were awake, we did not model the participant’s cognitive state, for example, using participant’s self-reported subjective assessments of their cognitive state during the RS measurement (Diaz et al. 2013). Since the cognitive state and the extent of its fluctuation during rest are difficult to establish for each individual, the high identification accuracy with *RS1* might have been attributable to highly motivated and instruction-compliant participants rather than the neural characteristics of the rest state. Therefore, the *Tap* and *Sequence* tasks provided verifiable within-subject examples of states that deviated from rest in order to assess the generality of RS-based inferences.

In a longitudinal setting, the classification problem of interest requires a decision between *NP*+ and *NP*-within the same individual. However, here *NP*+ was defined based on samples of RS activity from *other* individuals. This use of inter-individual differences provided a pragmatic means to simulate a diverse range of possible changes to an individual’s neurophysiology (Figure 1B) with the assumption that detecting true within-subject neurophysiological change would possibly be far more challenging. For example, in the *Sequence* task, the motor learning across the five days in our experiment involved neuroplastic changes (Wymbs et al. 2012; Wymbs and Grafton 2014; Bassett et al. 2015) and the accompanying changes in *SeqWait* and *SeqMov* over the duration of the experiment (Figure 8) could be considered as consequence of this learning-induced neuroplasticity. However, due to the unclear carryover effects of these plastic changes on *RS1* over this five day period, we instead used the *SeqWait* and *SeqMov* to simulate incidental cognitive-state variations (*NP-*) with high inter-day variance, where the neural dynamics on each day was a poor model of the dynamics on other days.

### 4.2. Reliability of identity inferences versus reliability of features

Numerous prior studies have investigated the inter-day similarity in RS activity within a test-retest framework (Bijsterbosch et al. 2017; Cox et al. 2018; Noble et al. 2019; Postema et al. 2019). In that framework, the focus is on evaluating whether a particular measure of RS activity on day *p* was reliably reproduced for RS activity on day *q* in the assumed <u>absence</u> of a true change. However, our focus is on the reliability of inferences in the assumed <u>presence</u> of true inter-day activity changes. This focus required differing considerations about how an individual’s unique identity was defined and represented as illustrated with an analogy to object recognition.

Consider images of the same object *X* from day 1 (test) and day 2 (retest) (Figure 11) represented by a list of filled pixel locations (i.e., features). With this representation, a simple measure of test-retest reliability is whether a pixel’s filled state on day 1 is a reliable predictor of its state on day 2. The scenario in Figure 11A is consistent with a *high* feature-level reliability as the majority of filled pixels on day 1 are also filled on day 2. However, this high reliability is misleading about the object’s unique identity. On day 1, object *X* can be readily distinguished from object *Y* based on a few critical pixels (circled). These critical pixels on object *X* are, however, unfilled on day 2. Thus, object *X* is not uniquely identifiable on day 2 as it is now confusable with object *Y*. Conversely, in the scenario shown in Figure 11B, a pixel-based test of reliability would indicate a *low* reliability due to the large number of filled pixels from day 1 that are unfilled on day 2. However, this low reliability is a limitation of how the object was represented (i.e., as a list of filled pixel locations relative to the main axes). If this representation included information about the relationships between the filled locations, then the object’s defining characteristics would be deemed as being reliably conserved on day 2, e.g., a rotation of the object *X* on day 2 would bring it into correspondence with its form on day 1.

**Figure 11:**
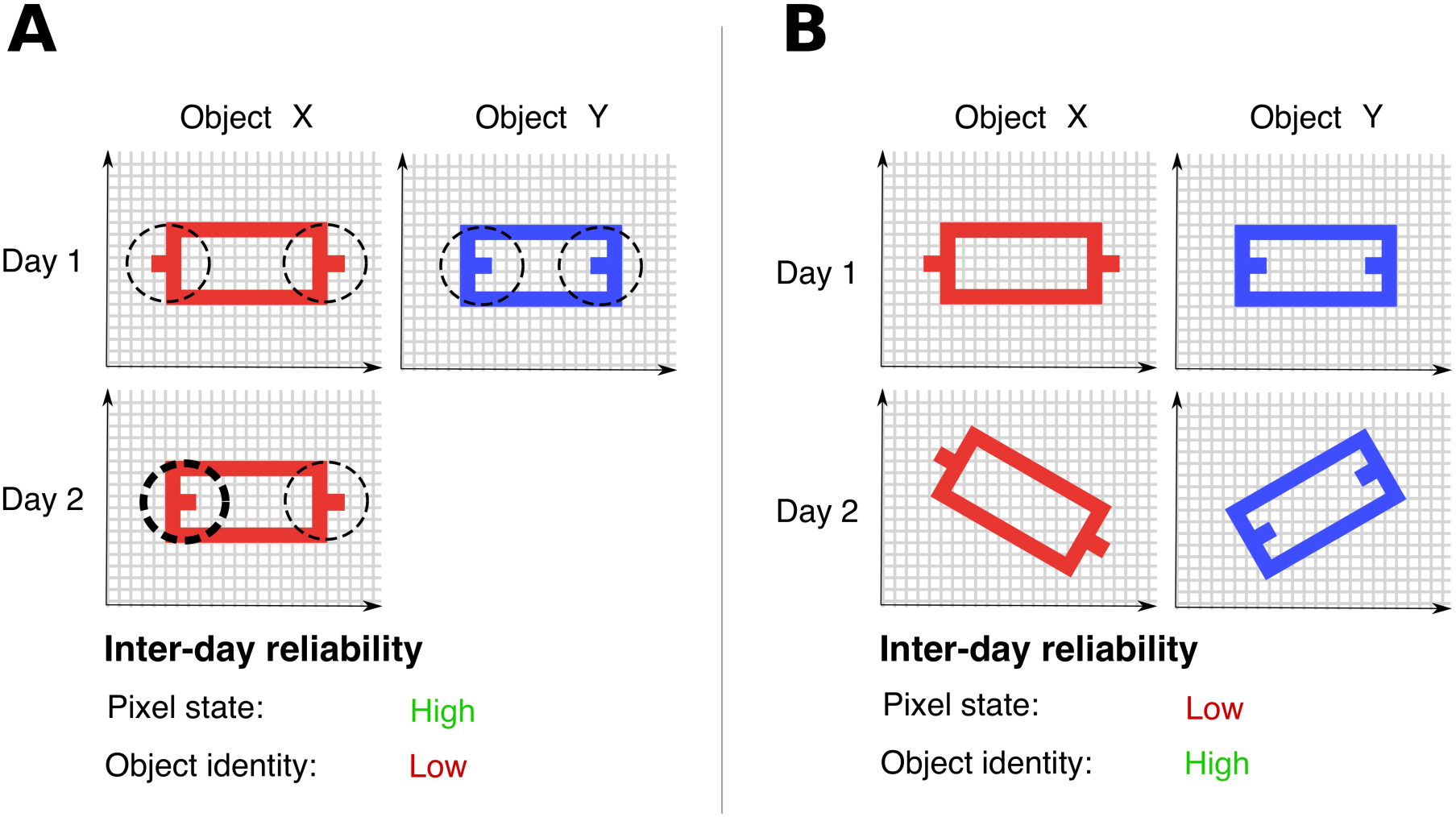
Test-retest reliability versus individual re-identification. **(A)** Objects *X* (red) and *Y* (blue) are uniquely defined by the configuration of filled and unfilled pixels. On day 1, (top row), the dotted circles indicates the critical pixels that distinguish *X* and *Y*. Most pixels of object *X* from day 1 are also filled on day 2 (lower row). However, pixels in the left dotted circle on day 2 differ in their state from day 1. Due to this difference on day 2, object *X* cannot be uniquely re-identified as being object *X* based on its form as it is now confusable with object *Y.* High inter-day reliability in pixel state does not imply the same for object identity. (**B**) The orientation of objects *X* on day 2 is rotated relative to its orientations on day 1. If this orientation is accounted for, then object *X* can be uniquely re-identified on day 2. However, when considering individual pixels, most of the filled pixels on day 1 are not on day 2. Low inter-day reliability in pixel state does not imply the same for object identity.

As demonstrated by this analogy, high test-retest reliability of individual features does not imply the reliability of the configuration of these features to enable individual identification and vice versa. This relationship between reliability and how an individual’s identity is defined and represented was a central consideration here.

Despite using an analogy of an individual’s configuration to a static object in the above example, the core variance model in our analysis involved an assumption about time and time-scales. Each same-day measurement was segmented into 2s non-overlapping epochs where each epoch was treated as a sample drawn from an underlying individual-specific distribution. The dynamic variability between samples was assumed to be an important characteristic of this individual-specific distribution. Cross-day/cross-task identification was predicated on whether training classifiers based on the inter-sample variability on short time-scales (i.e., between the samples acquired within seconds/minutes of each other on the same day) was a viable model for samples obtained on long time-scales, i.e., days apart (Figure 1).

Even though we do not use an explicit model of functional connectivity, the multivariate representations used to represent an individual’s decision-rule assumes a coupling between power values across distributed locations. A distinction is often drawn between static and dynamic connectivity based on how the neural time-series over the resting task is interpreted (Hutchison et al. 2013; Calhoun et al. 2014). Static connectivity refers to the extraction of a single measure (e.g., a graph) from the time-series. In contrast, dynamic connectivity is based on the view that resting state refers to a collection of states that dynamically vary at different time points. However, our approach and findings here are agnostic as to whether the inter-sample differences indicate variability around a characteristic mean value (i.e., static connectivity) or characteristic transitions between distinct states (i.e., dynamic connectivity). The relationship between the classifier-based multivariate representations to connectivity and distances measures (e.g., Valizadeh et al. 2019; Pani et al. 2020) is a key issue to be resolved by future studies.

### 4.3. Individual identification and longitudinal tracking

By using individual differences as a source of neurophysiological information, here the problem of distinguishing between *NP*+ and *NP*-was equivalent to the problem of individual identification with similarities to numerous studies that have, for example, sought to use RS-EEG as an individual-specific signature for biometric identification (Campisi and Rocca 2014; Gui et al. 2015; Valizadeh et al. 2019). However, our focus was not biometric identification or the important issues related to the neural basis of individual differences and trait-identification (Smit et al. 2005, 2006; Demuru et al. 2017; Finn et al. 2017; Gratton et al. 2018). Nevertheless, our findings are consistent with these prior studies in demonstrating the high distinctiveness of individual differences and its robust detectability even across days and tasks from two-second snapshots of the oscillatory power spectra at rest.

However, the inter-individual differences in cross-day identification with *RS1* both with and without aggregation (Figure 5, 6A, 6C) also demonstrated that resting activity was not the strict equivalent of a “fingerprint”, i.e., in being entirely immune to cognitive state or even whether a person is alive (Campisi and Rocca 2014). Even though individual identification was possible across tasks with high accuracy, the RS-based individual signatures were not completely independent of cognitive state. Large deviations from rest during *TapMov* and *SeqMov* reduced cross-task identification accuracy even though identification was above random chance. These accuracy reductions were due to cognitive state differences with *RS1* and not merely because *TapMov* and *SeqMov* conditions lacked identifiable signatures or had more movement-related artifacts (Figure 8D).

The use of an individual identification strategy involved certain tradeoffs. An individual’s identity was defined by analyzing differences to other individuals in the studied group. Therefore, the characteristics represented by an individual *S*_X_’s decision-rule could vary depending on the properties of the other individuals in the group. Rather than the number of individuals in the group, the key determinants of how *S*_X_ is represented would be the diversity and properties of the most-similar individuals (as illustrated by the confusion matrix and inter-individual clustering in Figure 5). Furthermore, identifying features that distinguish an individual from others would lead to the exclusion of features *shared* by all individuals. For example, in a study of the heritability of individual RS-connectivity properties with magnetoencephalography (MEG) (Demuru et al. 2017), the explicit removal of connectivity characteristics shared by all individuals in the group was found to significantly improve individual identification. However, down-weighting the role of shared features (explicitly or implicitly) has a tradeoff for tracking neural plasticity since changes to an individual’s neurophysiology on these shared features might go undetected.

### 4.4. What makes an individual configuration robust to changes in cognitive state?

Our primary findings are based on black-box statistical inferences, namely, the pattern of classification accuracies obtained with different training/test sets that were selected based on experimental variables (e.g., the effect of day, the conditions defining the cognitive state, and the feature set). Therefore, an important issue is whether these statistical regularities are consistent with a neurophysiological signature in RS-activity rather than a byproduct of other factors specific to our implementation.

The shape of the power spectrum in the frequency domain at rest has long been suggested as an important individual characteristic (Näpflin et al. 2007; Chiang et al. 2011; Bazanova and Vernon 2014). This shape has multiple peaks over an aperiodic background of 1/f noise. The specific frequencies at which these peaks occur, particularly in the α band and in the β band have been the topic of considerable investigation (van Albada and Robinson 2013; Voytek et al. 2015). Importantly, in different cognitive states, the changes to this spectrum are not arbitrary and primarily involve changes to the power at the peaks (as well as small shifts in the peak frequency) but without large changes to the 1/f background (Buzsáki et al. 2012; Haegens et al. 2014; Cole and Voytek 2017). Furthermore, Demuru and Fraschini (2020) found that this aperiodic background was highly individual-specific and allowed individuals to be identified with higher accuracy than the power in canonical frequency bands.

Therefore one possibility to explain our results is that an individual’s decision-rule implicitly represents the shape of their unique power spectrum. If this were the case then it would provide a plausible explanation for the observed high specificity despite cognitive state variation. In our feature representation, the power over the full power spectrum was averaged into five canonical bands. Therefore, capturing the individual shape of the spectrum and, for example, the approximate location of the α power peak would require a role for features representing the power in the θ, α and β_1_ bands. Indeed, it was these three bands that also showed the main consistencies in term of a few, high valued weights. The classical depiction of the power spectrum is from a particular channel. Our finding suggests that representation of the individual-specific power in the different bands were distributed over the scalp with a concentration in the fronto-central and occipital zones. Although the power spectra are similar across channels, any one channel is an incomplete representation of that individual’s characteristic power distribution. Consequently, it might lack the robustness to represent individual variability across days. By contrast, a decision-rule that combines each band’s best representation might have a greater robustness.

### 4.4. Outlook

In the current study, we assumed that individuals in the studied group did not undergo extensive plastic changes. If individual identification was not possible with longitudinal RS even with such a group of healthy individuals over a period of five days, then the merits of using RS as a tracking indicator would seem to require critical re-evaluation especially for tracking over longer periods of time and with populations where such neuroplastic changes would be expected. Prior studies have found changes to the power spectrum with aging (van Albada et al. 2010; Chiang et al. 2011; Voytek et al. 2015; Knyazeva et al. 2018), for example, age-related reductions in the frequencies of the alpha and beta band peaks. Voytek et al. (2015) suggest that such changes might indicate a change in the 1/f baseline possibly due to increased physiological noise with aging. Furthermore, systematic longitudinal changes in the power spectrum have been observed following stroke (Giaquinto et al. 1994; Saes et al. 2020). Thus, the application of this physiological signature to monitor longitudinal RS in clinical populations is an important future priority.

## Acknowledgments

This work was funded by the University of Cologne Emerging Groups Initiative (CONNECT group) implemented into the Institutional Strategy of the University of Cologne and the German Excellence Initiative; and by the Deutsche Forschungsgemeinschaft (DFG, German Research Foundation) – Project-ID 431549029 – SFB 1451. SD gratefully acknowledges support from the German Research Foundation (DA 1953/5-2). We thank Hannah Kirsten, Alexandra Kurganova, and members of the INM-3 for their valuable assistance in data acquisition.

## CRediT author statement

**Maximilian Hommelsen**: Conceptualization, Methodology, Software, Validation, Formal analysis, Investigation, Writing: Original Draft, Visualization. **Shivakumar Viswanathan**: Conceptualization, Methodology, Software, Validation, Formal analysis, Writing: Review & Editing, Visualization. **Silvia Daun:** Conceptualization, Writing: Review & Editing, Visualization, Supervision, Project administration, Funding acquisition.

## APPENDIX

**Table A.1:** Identification accuracies in different experimental states reported as Mean % (SD). All values were significantly above random chance (50%) (see Suppl. Table 1).

|  |  |  |  |  |
| --- | --- | --- | --- | --- |
| **States** | **Type** |  |  |  |
|  | **CV** | **1-day** | **2-day** | **3-day** |
| ***RS1***<br>(N=24) | 99.98<br>(0.04) | 92.10<br>(6.84) | 95.93<br>(3.63) | 97.39<br>(2.65) |
| ***RS1***<br>(N=18) | 99.98<br>(0.06) | 92.79<br>(6.76) | 96.61<br>(3.30) | 97.53<br>(2.51) |
| ***RS2***<br>(N=24) | 99.99<br>(0.04) | 91.58<br>(7.49) | 95.86<br>(4.18) | 96.99<br>(3.50) |
| ***TapWait***<br>(N=18) | 99.99<br>(0.02) | 92.58<br>(6.39) | 96.36<br>(3.42) | 97.60<br>(2.77) |
| ***SeqWait***<br>(N=18) | 99.99<br>(0.02) | 93.67<br>(7.35) | 97.12<br>(4.36) | 98.03<br>(3.80) |
| ***TapMov***<br>(N=18) | 99.94<br>(0.12) | 92.39<br>(6.72) | 96.12<br>(3.34) | 97.29<br>(2.41) |
| ***SeqMov***<br>(N=18) | 100.00<br>(0.00) | 93.47<br>(8.41) | 96.67<br>(4.64) | 97.95<br>(2.99) |

**Table A.2:** Identification accuracies for *RS1* with mono-band and mono-location feature subsets reported as Mean % (SD). All values were significantly above random chance (50%) (see Suppl. Table 2).

| **Table A.2:** Identification accuracies for *RSI* with mono-band and mono-location feature subsets reported as Mean % (SD). All values were significantly above random chance (50%) (see Suppl. Table 2). |  |  |  |  |
| --- | --- | --- | --- | --- |
| **Subset**<br>(*N*=24) | **Type** |  |  |  |
|  | **CV** | **1-day** | **2-day** | **3-day** |
| $B_{\delta}$ | 96.10<br>(2.54) | 64.66<br>(7.92) | 67.87<br>(8.12) | 70.12<br>(8.01) |
| $B_{\theta}$ | 97.63<br>(1.52) | 76.99<br>(7.69) | 81.76<br>(7.11) | 83.70<br>(6.94) |
| $B_{\alpha}$ | 98.51<br>(1.17) | 84.20<br>(7.74) | 88.38<br>(6.34) | 89.59<br>(5.67) |
| $B_{\beta 1}$ | 99.65<br>(0.57) | 81.41<br>(10.44) | 87.03<br>(9.03) | 88.92<br>(8.14) |
| $B_{\beta 2}$ | 99.74<br>(0.30) | 76.37<br>(9.98) | 83.22<br>(9.00) | 86.66<br>(7.96) |
| $L_F$ | 98.01<br>(1.80) | 82.68<br>(8.89) | 87.30<br>(7.45) | 88.87<br>(6.56) |
| $L_{FC}$ | 98.54<br>(1.55) | 86.93<br>(9.38) | 90.39<br>(7.45) | 91.76<br>(6.12) |
| $L_{CP}$ | 97.94<br>(1.78) | 85.28<br>(8.57) | 89.43<br>(7.30) | 90.37<br>(6.52) |
| $L_{PO}$ | 97.96<br>(2.02) | 81.02<br>(8.32) | 86.47<br>(7.35) | 87.97<br>(7.06) |

**Table A.3:** Classification accuracy of *RS1* vs task state (binary, within-subject) reported as Mean % (SD). All values were significantly above random chance (50%) (see Suppl. Table 3).

| **Table A.3:** Classification accuracy of *RSI* vs task state (binary, within-subject)<br>reported as Mean % (SD). All values were significantly above random chance (50%)<br>(see Suppl. Table 3). |  |  |  |  |
| --- | --- | --- | --- | --- |
| *RSI* vs<br>(*N*=18) | Type |  |  |  |
|  | CV | 1-day | 2-day | 3-day |
| *TapWait* | 88.35<br>(5.66) | 62.91<br>(6.44) | 66.26<br>(8.69) | 67.28<br>(9.11) |
| *SeqWait* | 95.12<br>(3.74) | 67.79<br>(8.53) | 73.05<br>(11.01) | 74.86<br>(11.37) |
| *TapMov* | 93.56<br>(4.12) | 79.04<br>(7.17) | 82.75<br>(5.99) | 84.18<br>(5.92) |
| *SeqMov* | 97.81<br>(1.76) | 88.77<br>(5.21) | 92.81<br>(3.43) | 93.32<br>(3.77) |

**Table A.4:** Accuracy of cross-task ^RS1^I_p_ → ^X^I_q_ identification reported as Mean % (SD). All values were significantly above random chance (50%) (see Suppl. Table 4).

| **Table A.4:** Accuracy of cross-task $^{RSI}I_p \rightarrow ^XI_q$ identification reported<br>as Mean % (SD). All values were significantly above random chance<br>(50%) (see Suppl. Table 4). | | | |
| --- | --- | --- | --- |
| Test states<br>(*N* = 18) | Type |  |  |
|  | 1-day | 2-day | 3-day |
| *TapWait* | 91.90<br>(6.46) | 95.84<br>(3.30) | 96.90<br>(2.44) |
| *SeqWait* | 90.81<br>(7.09) | 94.95<br>(4.39) | 96.09<br>(3.40) |
| *TapMov* | 88.79<br>(7.57) | 93.02<br>(5.49) | 94.01<br>(4.51) |
| *SeqMov* | 83.85<br>(10.35) | 88.39<br>(9.28) | 90.03<br>(8.87) |

## SUPPLEMENTARY MATERIAL

**Supplementary Table 1:**
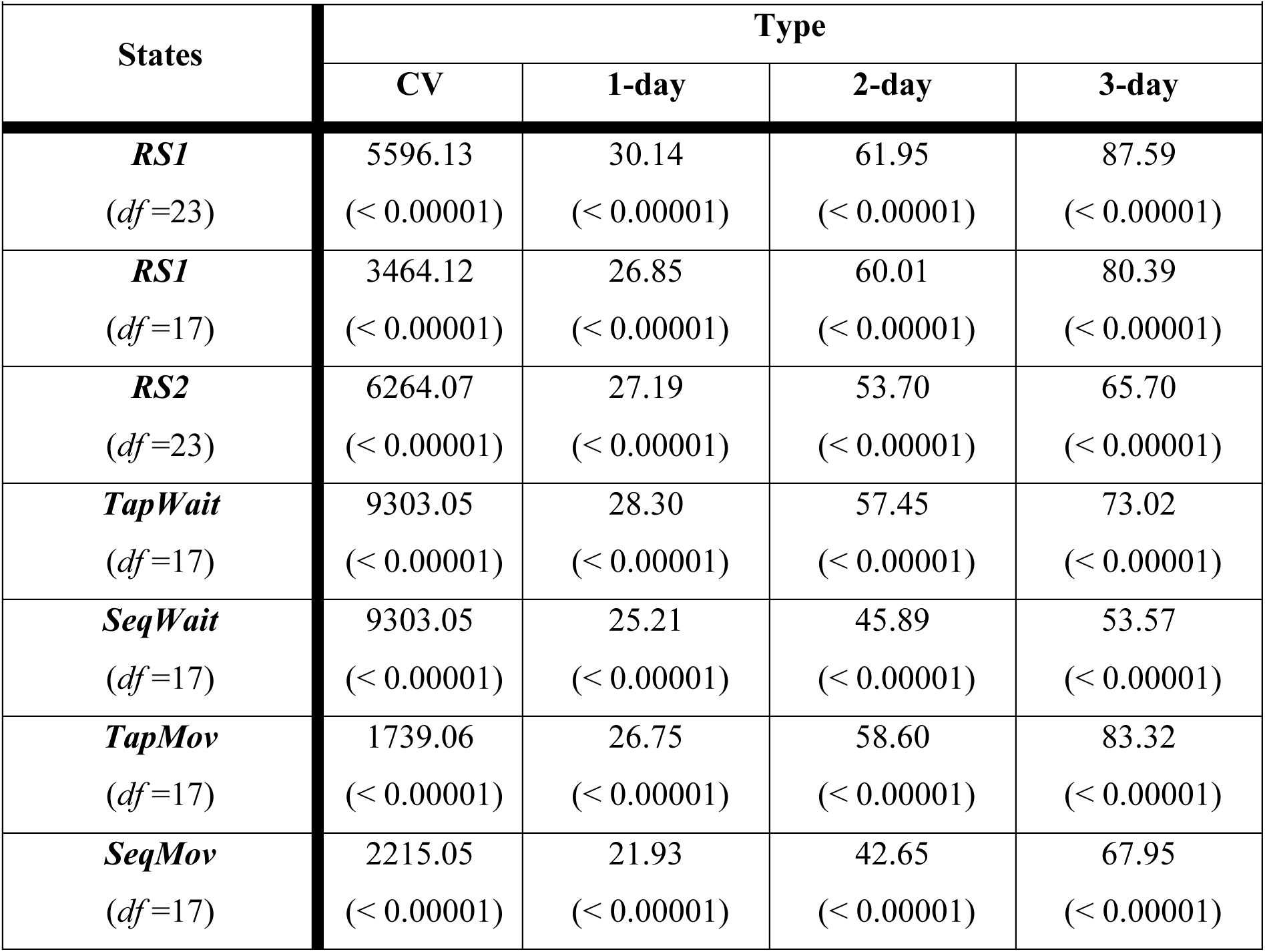
One-sample t-tests of mean identification accuracy in different states (see Table A.1) vs random chance (50%) reported as t-value (p-value).

|  |  |  |  |  |
| --- | --- | --- | --- | --- |
| **States** | **Type** |  |  |  |
|  | **CV** | **1-day** | **2-day** | **3-day** |
| ***RS1***<br>(*df*=23) | 5596.13<br>($< 0.00001$) | 30.14<br>($< 0.00001$) | 61.95<br>($< 0.00001$) | 87.59<br>($< 0.00001$) |
| ***RS1***<br>(*df*=17) | 3464.12<br>($< 0.00001$) | 26.85<br>($< 0.00001$) | 60.01<br>($< 0.00001$) | 80.39<br>($< 0.00001$) |
| ***RS2***<br>(*df*=23) | 6264.07<br>($< 0.00001$) | 27.19<br>($< 0.00001$) | 53.70<br>($< 0.00001$) | 65.70<br>($< 0.00001$) |
| ***TapWait***<br>(*df*=17) | 9303.05<br>($< 0.00001$) | 28.30<br>($< 0.00001$) | 57.45<br>($< 0.00001$) | 73.02<br>($< 0.00001$) |
| ***SeqWait***<br>(*df*=17) | 9303.05<br>($< 0.00001$) | 25.21<br>($< 0.00001$) | 45.89<br>($< 0.00001$) | 53.57<br>($< 0.00001$) |
| ***TapMov***<br>(*df*=17) | 1739.06<br>($< 0.00001$) | 26.75<br>($< 0.00001$) | 58.60<br>($< 0.00001$) | 83.32<br>($< 0.00001$) |
| ***SeqMov***<br>(*df*=17) | 2215.05<br>($< 0.00001$) | 21.93<br>($< 0.00001$) | 42.65<br>($< 0.00001$) | 67.95<br>($< 0.00001$) |

**Supplementary table 2:** One-sample t-tests of mean identification accuracy for *RS1* with mono-band and mono-location feature subsets (see Table A.2) vs random chance (50%) reported as t-value (p-value).

| **Supplementary table 2:** One-sample t-tests of mean identification accuracy for *RSI* with mono-band and mono-location feature subsets (see Table A.2) vs random chance (50%) reported as t-value (p-value). |  |  |  |  |
| --- | --- | --- | --- | --- |
| **Subset**<br>(*df*=23) | **Type** |  |  |  |
|  | **CV** | **1-day** | **2-day** | **3-day** |
| $B_{\delta}$ | 88.62<br>($< 0.00001$) | 9.07<br>($< 0.00001$) | 10.79<br>($< 0.00001$) | 12.29<br>($< 0.00001$) |
| $B_{\theta}$ | 151.87<br>($< 0.00001$) | 17.20<br>($< 0.00001$) | 21.88<br>($< 0.00001$) | 23.79<br>($< 0.00001$) |
| $B_{\alpha}$ | 202.00<br>($< 0.00001$) | 21.64<br>($< 0.00001$) | 29.69<br>($< 0.00001$) | 34.22<br>($< 0.00001$) |
| $B_{\beta 1}$ | 415.09<br>($< 0.00001$) | 14.73<br>($< 0.00001$) | 20.09<br>($< 0.00001$) | 23.42<br>($< 0.00001$) |
| $B_{\beta 2}$ | 703.68<br>($< 0.00001$) | 12.95<br>($< 0.00001$) | 18.10<br>($< 0.00001$) | 22.61<br>($< 0.00001$) |
| $L_F$ | 131.76<br>($< 0.00001$) | 18.00<br>($< 0.00001$) | 24.50<br>($< 0.00001$) | 29.01<br>($< 0.00001$) |
| $L_{FC}$ | 154.96<br>($< 0.00001$) | 19.29<br>($< 0.00001$) | 26.54<br>($< 0.00001$) | 33.46<br>($< 0.00001$) |
| $L_{CP}$ | 132.33<br>($< 0.00001$) | 20.15<br>($< 0.00001$) | 26.47<br>($< 0.00001$) | 30.29<br>($< 0.00001$) |
| $L_{PO}$ | 116.20<br>($< 0.00001$) | 18.26<br>($< 0.00001$) | 24.30<br>($< 0.00001$) | 26.33<br>($< 0.00001$) |

**Supplementary table 3:** One-sample t-tests of classification accuracy of *RS1* vs task state (see Table A.3) against random chance (50%) reported as t-value (p-value).

| **Supplementary table 3:** One-sample t-tests of classification accuracy of *RSI* vs task state (see Table A.3) against random chance (50%) reported as t-value (p-value). |  |  |  |  |
| --- | --- | --- | --- | --- |
| ***RSI* vs**<br>(*df*=17) | **Type** |  |  |  |
|  | **CV** | **1-day** | **2-day** | **3-day** |
| ***TapWait*** | 28.74<br>($< 0.00001$) | 8.51<br>($< 0.00001$) | 7.94<br>($< 0.00001$) | 8.05<br>($< 0.00001$) |
| ***SeqWait*** | 50.81<br>($< 0.00001$) | 8.85<br>($< 0.00001$) | 8.88<br>($< 0.00001$) | 9.28<br>($< 0.00001$) |
| ***TapMov*** | 45.23<br>($< 0.00001$) | 17.19<br>($< 0.00001$) | 23.18<br>($< 0.00001$) | 24.49<br>($< 0.00001$) |
| ***SeqMov*** | 119.60<br>($< 0.00001$) | 31.55<br>($< 0.00001$) | 52.90<br>($< 0.00001$) | 48.70<br>($< 0.00001$) |

**Supplementary table 4:** One-sample t-tests of cross-task ^RS1^**I**_p_ → ^X^**I**_q_ identification accuracy (see Table A.4) against random chance (50%) reported as t-value (p-value).

| **Supplementary table 4:** One-sample t-tests of cross-task $^{RSI}I_p \rightarrow ^XI_q$ identification accuracy (see Table A.4) against random chance (50%) reported as t-value (p-value). | | | |
| --- | --- | --- | --- |
| **Test states**<br>(*df* = 17) | **Type** |  |  |
|  | **1-day** | **2-day** | **3-day** |
| ***TapWait*** | 27.52<br>($< 0.00001$) | 58.90<br>($< 0.00001$) | 81.63<br>($< 0.00001$) |
| ***SeqWait*** | 24.43<br>($< 0.00001$) | 43.39<br>($< 0.00001$) | 57.55<br>($< 0.00001$) |
| ***TapMov*** | 21.73<br>($< 0.00001$) | 33.23<br>($< 0.00001$) | 41.42<br>($< 0.00001$) |
| ***SeqMov*** | 13.88<br>($< 0.00001$) | 17.55<br>($< 0.00001$) | 19.14<br>($< 0.00001$) |

**Supplementary Figure 1:**
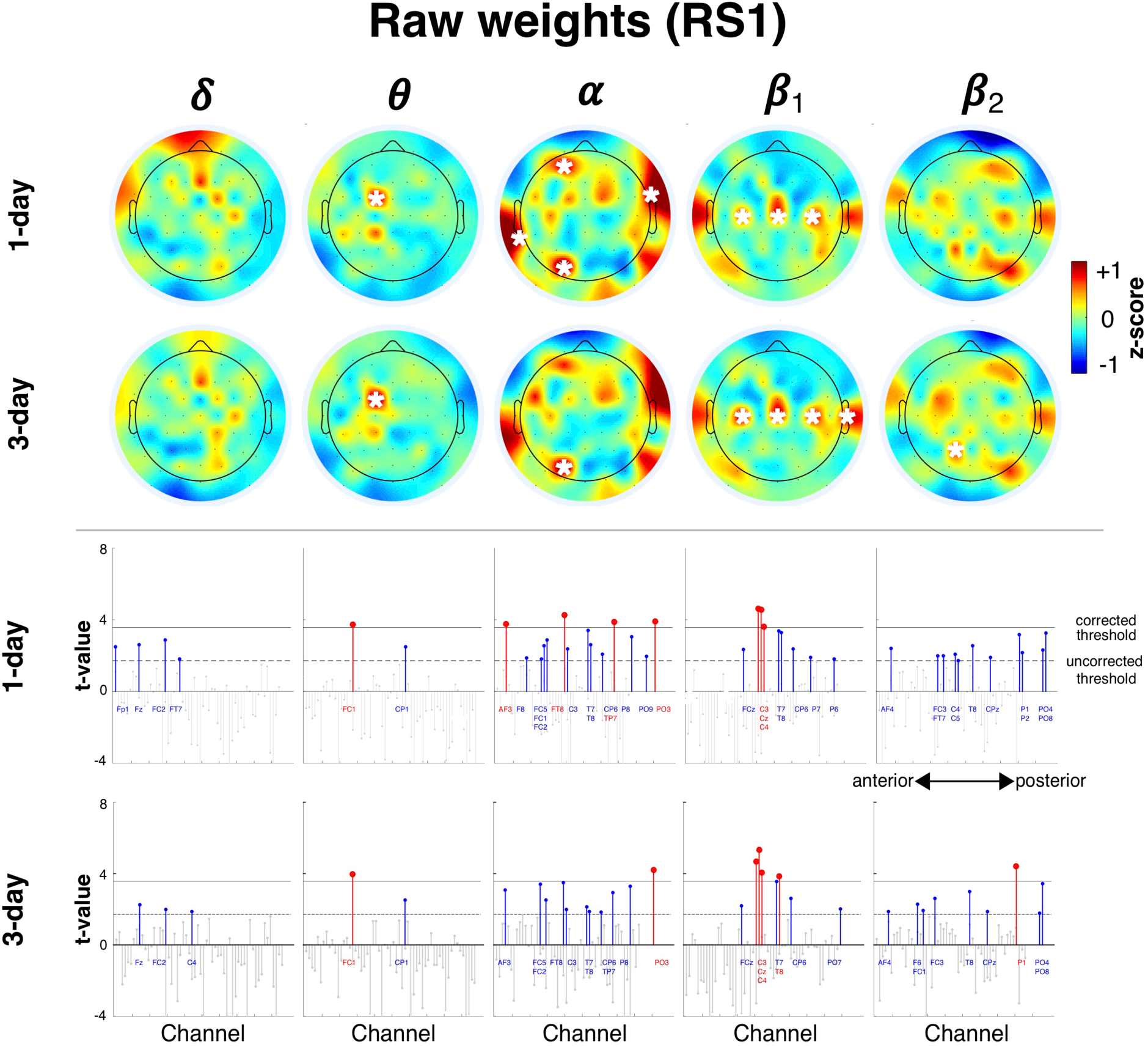
High-consistency features (raw weights). Spatial distribution of high-consistency raw weights for frequency bands of full feature set (z-scored across all features) and their aggregation-related changes (1-day, 3-day). Mean weights in each scalp map that were significantly greater than zero are indicated with a white asterisk (*p* < 0.05/61). Lower two rows show *t*-values for the features corresponding to the upper rows. Channels have an anterior-to-posterior ordering (x-axis). Red stems indicate channels with t-values higher than the corrected threshold (*p* < 0.05/61, horizontal black line) while blue stems show channels that only pass uncorrected thresholds (*p* <0.05, dotted horizontal line). Colored channel labels are grouped from top-to-bottom for visibility and correspond to stems from left to right.

### Supplementary Figure 2: Concentration of high-relevance features in *B_α_* and *L*_FC_

The feature weights were used to assess whether differences in cross-day accuracies between the mono-band and mono-location subsets were an indicator of their relevance in the full feature set. For example, if subset *S_x_* in isolation had a higher cross-day accuracy than *S_y_* then *S_x_* might have a larger concentration of high-weighted features than *S_y_* as part of the full feature-set. To test this simplistic prediction, we evaluated the relative concentration of features with large weights (specified by percentile) in the different mono-band (Suppl. Figure 2A) and mono-location subsets (Suppl. Figure 2B).

**Supplementary Figure 2:**
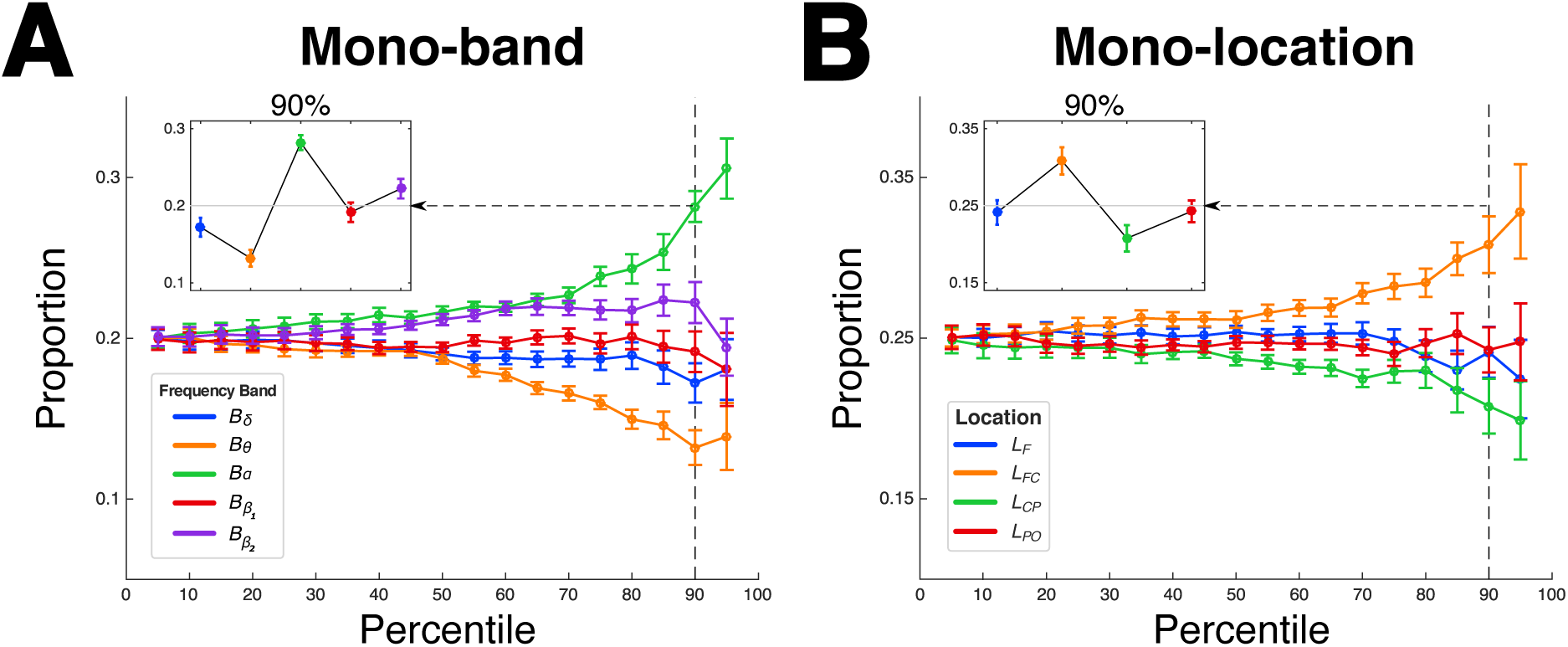
High-valued weights. Mean proportions of high-valued (absolute) weights in the mono-band subsets (panel **A**) and mono-location subsets (panel **B**) at increasing percentiles (x-axis). Proportions (y-axis) are shown as deviations from an equal distribution across subsets (1/5 = 0.2, upper panel; and 1/4 = 0.25; lower panel). Insets show weight distribution in the 90^th^ percentile (vertical dotted line). Error bars: Within-subject s.e.m

The distribution of high-valued weights in the mono-band/location subsets was estimated as follows. For each individual, the absolute weights of all features were first sorted. The weights in the *k^t^*^h^ percentile of the sorted weights were then identified and the relative proportion of these selected high-valued weights contained in each of the mono-band sets was then calculated. This procedure was repeated for different values of *k.* These proportions were separately calculated for the mono-location sets. Since the temporal channels were not included in any of the mono-location subsets, these channels were excluded for the proportion calculations.

The relative proportion of high-weighted features in the different mono-band subsets diverged at higher percentiles [ANOVA, Band {*B_δ_, B_θ_, B_α_, B_β1_, B_β2_*} x Percentile {5%, 10%…95%}, Band*Perc: F_72, 1656_ = 5.08, p < 0.00001; Band: F_4, 92_ = 7.26, p = 0.00003; Perc: F_18, 414_ < 0.001, p =1]. *B_α_* contained the largest proportion of high-valued weights in the 90^th^ percentile (inset, panel A). This ordering was qualitatively similar to the cross-day accuracies for the mono-band feature sets (Figure 6A), where *B_α_* had the highest mean cross-day accuracy. However, the proportion of high weights in *B_δ_* was comparable to the other subsets despite having a lower cross-day accuracy than the other subsets. The relative proportion of high-weighted features in the mono-location subsets also diverged at higher percentiles [ANOVA, Location: {*L*_F_, *L*_FC_, *L*_CP_, *L*_PO_} x Percentile {5%, 10%…95%}, Location*Perc: F_54, 1242_ = 2.00, p = 0.00003; Location: F_3, 69_ = 3.71, p = 0.015; Type: F_18, 414_ < 0.001, p = 1]. The fronto-central *L*_FC_ subset contained a higher proportion of high-valued weights in the 90^th^ percentile than *L*_CP_ that was immediately posterior to *L*_FC_ (inset, panel B). By contrast, the mean cross-day accuracies for *L*_FC_ and *L*_CP_ in isolation were comparably similar (Figure 6C).

The concentration of high-valued weights in the different feature subsets was not a simple indicator of how these feature sets might contribute to high cross-day accuracy. However, it revealed consistencies in the distribution of relevant features across individuals, notably in *B_α_* and *L*_FC_.

**Supplementary Figure 3:**
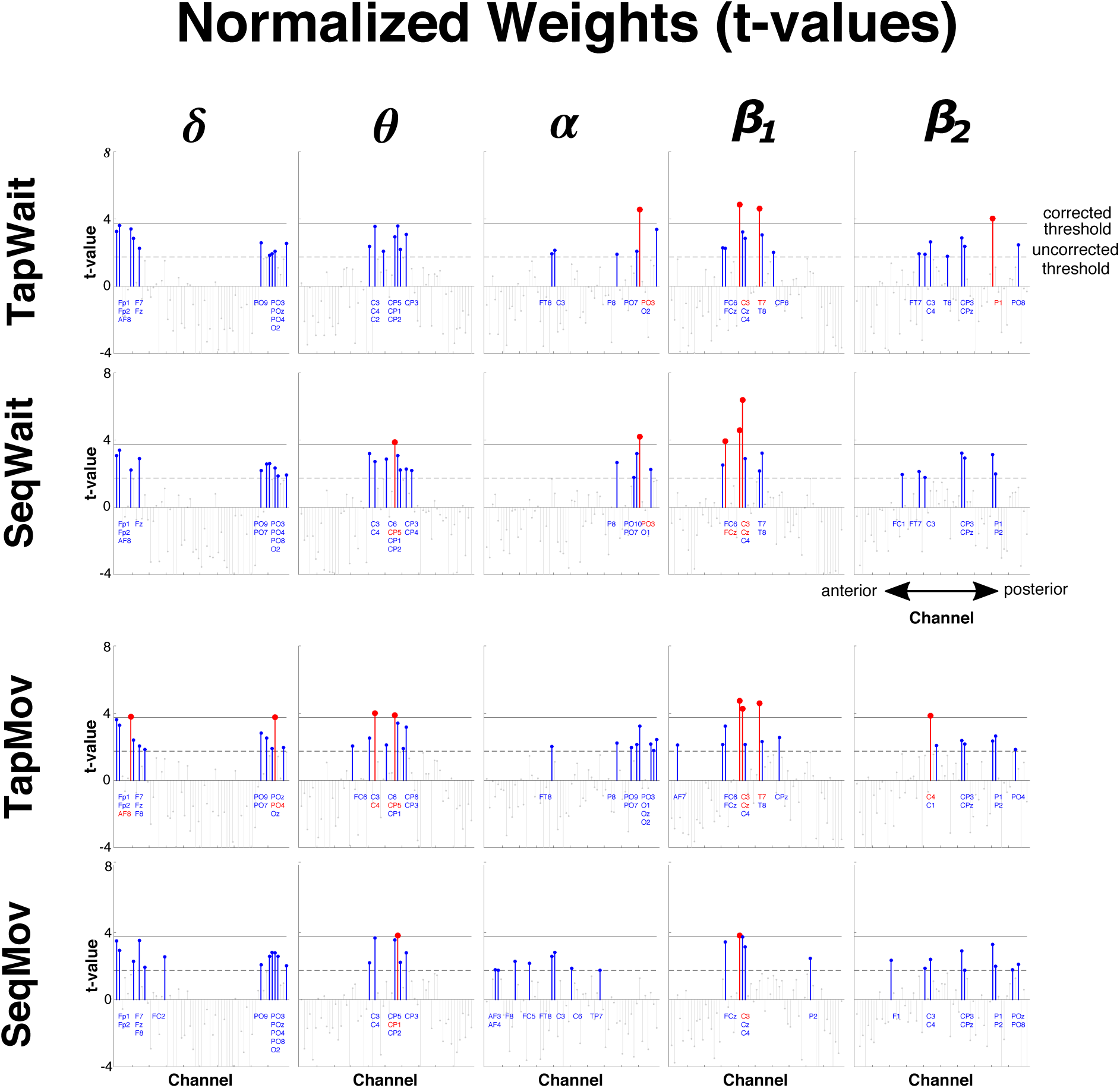
Stem plot of t-values of high-consistency weights (1-day) in task-states per frequency band from Figure 9A. All labeling conventions are as in Suppl. Figure 1.

